# PixelPaws: a low-cost, open-source integrated platform for automated scoring of mouse behaviors

**DOI:** 10.64898/2026.09.19.752787

**Authors:** Richard A. Slivicki, Hanyun Wang, Juliet M. Mwirigi, Kayla Kucz, Hunter Hua, Meaghan C. Creed, Robert W. Gereau

**Author notes:** Corresponding author: Richard A. Slivicki, Washington University Pain Center, Department of Anesthesiology, Washington University School of Medicine, St. Louis, MO.

## Abstract

Behavioral scoring is central to preclinical pain research but is typically performed manually, which is inherently low-throughput and subject to variability between observers. Automated alternatives exist, but cost and programming expertise limit adoption beyond the laboratories that build them. Here we describe PixelPaws, an open-source platform combining a near-infrared imaging enclosure built from consumer parts for under $300, multi-camera acquisition software, a bundled pose-estimation network, and eight classifiers in a graphical user interface. We evaluated the platform across standard assays of pain, itch, and opioid withdrawal. Licking, scratching, and jumping classifiers were validated against trained human observers by leave-one-session-out evaluation. In the formalin assay, classifier-scored licking followed the expected biphasic time course and was reduced dose-dependently by oxycodone, with a phase 2 potency estimate comparable to blinded manual scoring. Chloroquine roughly tripled scratching, and naloxone precipitated a burst of jumping in oxycodone-treated mice. From the same recordings we derived a set of paw contour metrics. Contour intensity fell after formalin and tracked the oxycodone dose series, and contour area shifted toward symmetry after oxycodone in nerve-injured mice. Behavioral sequencing revealed distinct changes in the transitions between behaviors after naloxone-precipitated withdrawal and formalin administration. Oxycodone suppressed formalin-evoked licking without restoring behavioral sequencing toward the vehicle state. These data demonstrate that an integrated, low-cost, open-source platform can make automated behavioral phenotyping accessible without requiring users to build a custom computational pipeline.

## 1. Introduction

Behaviors such as licking, scratching, and grooming are central readouts in preclinical pain and neuropharmacology research, yet they are often manually scored. Scoring one behavior in one 30 min recording takes 50 to 150 min of observer time, so a six-behavior analysis of a 60-subject experiment can require up to 900 h [29]. Manual scoring is also observer-dependent, and inter-rater variability, observer bias, and observer drift are recognized problems that limit objectivity and reproducibility [24,40]. Together, these limitations motivate automated, objective, and scalable measures of behaviors that can support large pharmacological cohorts and dose-response designs.

Deep learning-based pose estimation has made such measures possible. DeepLabCut and SLEAP track body-part keypoints in freely moving animals [27,32], and those keypoints can be converted into behavioral labels with downstream classifiers. Near-infrared (NIR) illumination of the plantar surface has attracted recent interest because it yields high-contrast images of the paws, the focus of many pain and itch research assays. Brightness increases as a paw presses against the floor, which supplies a contact cue that pose alone does not capture and isolates paw-specific quantities: light intensity and contact area vary with paw-floor contact and have been used as optical proxies for weight bearing [43]. BAREfoot established that DeepLabCut pose combined with NIR pixel-brightness features and a gradient-boosted classifier scores flinching, licking, and grooming at human level [4].

Despite this progress, adoption outside the laboratories that build these tools remains limited. Most behavioral neuroscience laboratories lack programming or deep learning expertise and face a high entry barrier to implementing published machine learning approaches [40].

Graphical packages such as JAABA and SimBA lower that barrier for classifier training [18,21], but they address analysis alone and leave hardware and acquisition to the user. Commercial all-in-one systems remove both barriers but are proprietary and cost prohibitive for many laboratories [5,6].

We therefore set out to build an integrated platform a laboratory could assemble and operate without programming experience. Here we describe PixelPaws, which combines in a single system a low-cost NIR enclosure built from consumer parts for under $300, multi-camera acquisition software that records four enclosures in parallel from one computer, a bundled pretrained DeepLabCut pose network, and supervised classifiers that score user-defined behaviors, all driven from a graphical user interface. We validated classifiers for hind-paw licking, scratching, and jumping against trained human observers, and applied them to formalin-evoked licking and its dose-dependent reversal by oxycodone, to spared nerve injury, to chloroquine-induced itch, and to naloxone-precipitated opioid withdrawal. We further use the same recordings to derive paw-level contour metrics and quantify changes in the sequencing of behavior. By packaging hardware, acquisition, pose estimation, and classification together and releasing them open source, PixelPaws is intended to make automated behavioral scoring practical for laboratories that do not otherwise have dedicated computational support.

## 2. Materials and methods

### 2.1. Animals

C57BL/6J mice were bred in house from breeders originally obtained from The Jackson Laboratory (Bar Harbor, ME) and were 8 to 12 weeks of age at the start of testing, with the exception of the spared nerve injury cohort described below. Mice were group housed 5 per cage on a 12:12 light-dark cycle with lights on at 0600, with food and water available ad libitum, and were tested during the light cycle. Group sizes are given in the figure legends. Both sexes were included in the formalin, oxycodone dose-response, and rimonabant cohorts; the spared nerve injury cohort was male only, and the chloroquine and naloxone-precipitated withdrawal cohorts female only. In the mixed-sex cohorts, male and female mice were recorded on separate days. Sex was therefore not compared explicitly, and data are collapsed across sex as the study was not powered to detect sex differences. Single-sex cohorts reflected the availability of animals when each cohort was run rather than a biological hypothesis; the validation endpoints, agreement between classifier and human scoring, are not expected to depend on sex, and both sexes are represented across the battery as a whole. All procedures were approved by the Institutional Animal Care and Use Committee at Washington University in St. Louis and were conducted in accordance with the guidelines of the International Association for the Study of Pain.

### 2.2. Drug treatments and behavioral assays

All subcutaneously (s.c.) administered drugs were dissolved in sterile saline and given in a volume of 5 ml/kg. Rimonabant was given intraperitoneally (i.p.) in a DMSO-based vehicle, as described below. Mice were habituated to the recording enclosure for 30 min before video acquisition began, and the floor was cleaned between recordings.

### Formalin

Mice received an intraplantar injection of 2% formalin (Sigma-Aldrich, St. Louis, MO; #252549, 37% formaldehyde solution, diluted to 2% v/v in sterile saline, equivalent to approximately 0.74% formaldehyde), or saline vehicle into the left hind paw in a volume of 10 µl, and were recorded for 60 min immediately afterward. The response to intraplantar formalin is biphasic, comprising an acute first phase attributed to direct nociceptor activation and, after a brief quiescent interval, a prolonged second phase attributed to inflammation and central sensitization [15,38]. Licking was analyzed over the full session and summarized as phase 1 (0 to 10 min) and phase 2 (10 to 60 min).

### Oxycodone dose-response against formalin

Mice received oxycodone (Sigma-Aldrich #1485205; 1, 3, or 10 mg/kg, s.c.) or vehicle 5 min before intraplantar formalin, and mice were recorded for 60 min post-formalin injection.

### Spared nerve injury

Spared nerve injury was produced at approximately 8 weeks of age by ligation and transection of the tibial and common peroneal branches of the sciatic nerve, leaving the sural branch intact [14]. Behavioral testing was performed approximately 8 weeks after surgery, when mice were approximately 16 weeks of age, so testing falls in the maintained phase of the injury rather than during its development. Mice were recorded at baseline and again for 30 min post-drug following oxycodone (3 mg/kg, s.c.) or vehicle.

### Chloroquine-induced itch

The nape of the neck was shaved 1 day before testing, and mice received chloroquine (Sigma-Aldrich #C6628; 200 µg in 50 µl, s.c., into the nape) or an equal volume of saline, as previously described by our group [28,33,39]. Mice were recorded for 30 min post-drug.

### Rimonabant-evoked scratching

Rimonabant (Cayman Chemical, Ann Arbor, MI; #9000484) was dissolved in DMSO at 25 mg/ml to create stocks that were frozen until use. On the day of the experiment, stocks were diluted with DMSO to a final concentration of 20% DMSO, then 95% ethanol was added, followed by Tween 80 and finally saline, giving a vehicle of 20% DMSO, 8% ethanol, 8% Tween 80 and 64% saline, as described previously by our group [37]. Mice received rimonabant (1, 3 or 10 mg/kg, i.p., 5 ml/kg) or vehicle and were recorded for 60 min.

### Naloxone-precipitated withdrawal

Mice received a single dose of oxycodone (9 mg/kg, s.c.) and withdrawal was precipitated 2 h later with naloxone (Cayman Chemical #15594; 3 mg/kg, s.c.). The agonist to antagonist interval and the subcutaneous route follow established precipitated-withdrawal paradigms in C57BL/6J mice, in which morphine is followed 2 h later by naloxone [8,9,26]. Mice were recorded for 30 min before and after naloxone administration.

Mice were assigned to treatment groups at random. Experimenters performing injections and behavioral testing, and the annotators who scored video, were blinded to treatment; the classifiers score every recording identically and carry no information about group assignment.

### 2.3. Apparatus and video acquisition

The recording apparatus (Figure 1A) consisted of a 180 x 180 mm chamber with 110 mm walls, formed by opaque colored-acrylic panels standing on a transparent cast-acrylic floor (Chemcast clear acrylic, TAP Plastics, San Leandro, CA; approximately 92% transmission). The chamber sat within a light-tight filming box of 200 x 200 x 150 mm and 10 mm wall thickness. The chamber was illuminated from within by an 850 nm LED strip. Mice were imaged from directly below through the clear floor; 850 nm lies beyond the range of mouse cone sensitivity and is therefore effectively invisible to the mouse [20]. Each enclosure was built from a USB camera (See3CAM_CU27, e-con Systems, Chennai, India; Sony STARVIS IMX462 sensor) with an IR-corrected M12 varifocal lens (CV-2812-3MP, Marshall Electronics, Torrance, CA; 2.8 to 12 mm, f/1.4), an 850 nm SMD5050 LED strip (12 V DC, 60 LEDs/m, 14.4 W/m) driven through a regulated DC buck converter (DROK, Guangzhou, China; 5.3 to 32 V input, 12 A), cut acrylic and 3D-printed parts, for a total of $281.74 in materials. The complete bill of materials, with suppliers and current unit costs, is provided in Table 1.

**Figure 1.**
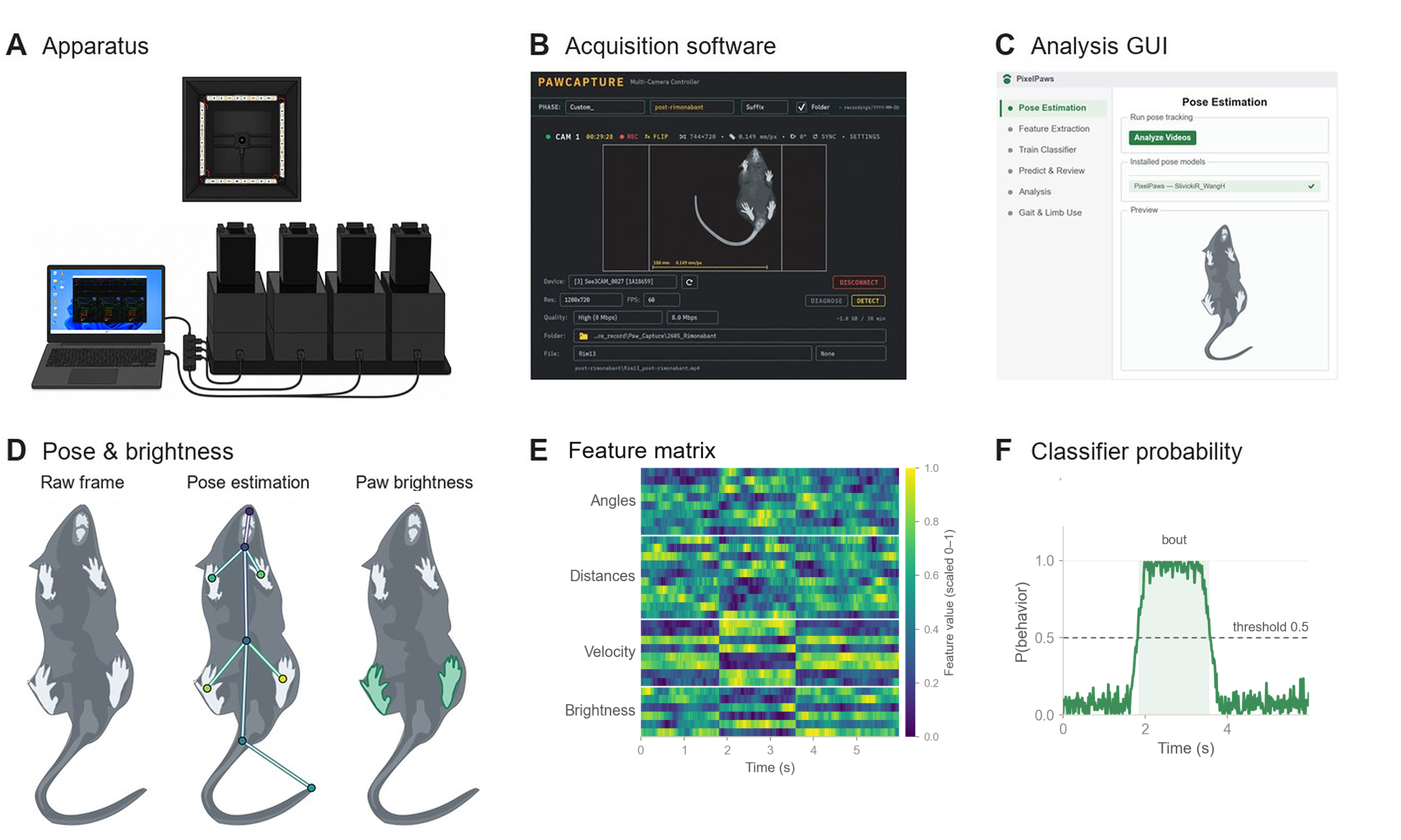
Overview of an open-source integrated platform for automated scoring of mouse behavior. Mice were recorded from below through a transparent floor under infrared illumination in parallel behavioral chambers controlled from a single laptop (A). Acquisition is handled by custom multi-camera software that records synchronized video with calibrated spatial scaling (B), and all downstream processing is performed in an integrated analysis application covering pose estimation, feature extraction, classifier training, prediction review and gait analysis (C). For each frame, markerless pose estimation predicts nine body parts and paw brightness is extracted from the region surrounding each paw keypoint, the latter reflecting contact with the acrylic (D). Pose and brightness are converted into a per-frame feature matrix comprising joint angles, inter-part distances, velocities, paw brightness and optical flow (E), which a supervised classifier converts into a per-frame probability for each behavior; contiguous frames exceeding threshold are merged into bouts (F).

The buck converter was added after the current-limiting resistors on the LED strip were found to warm the floor. Reducing the supply voltage removed the heating at the cost of dimmer illumination, which was compensated by raising camera exposure. Both configurations were represented in the recordings reported here and the behavioral readout did not differ between them (Figure S2), so either configuration can be used.

The camera connected over USB as a standard UVC device requiring no proprietary drivers. Recordings were acquired with PawCapture (v0.6.0), a multi-camera application distributed along with PixelPaws that drives several enclosures in parallel from a single Windows host, records each camera to its own hardware-encoded MP4 (NVENC, Quick Sync or AMF where available, with a CPU x264 fallback) at 720p and 60 fps (UYVY). Each session also wrote a single JSON manifest covering all cameras in that session, which the analysis tool reads to associate each file with its camera, phase and calibration. Per-camera settings are stored as reusable profiles and recordings can be tagged by experimental phase. Any UVC-compliant camera that is sensitive at 850 nm and lacks an infrared-cut filter can be used with PawCapture, although the bundled models were trained and validated only on recordings from the reference camera.

### 2.4. Pose estimation and feature extraction

PixelPaws produces frame-by-frame behavioral labels through a three-stage pipeline: pose estimation, feature extraction, and classification. Mice were recorded at 60 fps and pose was estimated with DeepLabCut [27] using a ResNet-50 backbone to predict nine body parts: all four paws, snout, neck, centroid, tail base, and tail end. The bundled network was trained on 2,482 manually labeled frames from 65 mice across 79 sessions spanning the formalin, oxycodone dose-response, spared nerve injury, chloroquine, rimonabant and naloxone-precipitated withdrawal cohorts, reaching a mean test error of 10.5 px, or 4.7 px for keypoints detected above a likelihood of 0.6, computed at the acquisition resolution of 1280 x 720. The network was trained on these recordings rather than initialized from a species-level pretrained pose model [42]. The trained network ships with the platform (section 2.11).

A feature matrix was then constructed from the pose estimates, consisting of inter-part distances, joint angles, velocities and accelerations, and their lags, together with keypoint visibility and height features and paw brightness sampled in a fixed window around each hind-paw and snout keypoint. Every pose-derived feature was computed between keypoints rather than relative to the chamber, so the feature matrix was invariant to the mouse’s position and orientation within the enclosure.

### 2.5. Behavior classifiers

BORIS event logs [17] were used to manually annotate behaviors. A built-in converter expanded each event log into a per-frame binary label within the observation window, 1 where the behavior was present and 0 where it was absent.

Behaviors were annotated to the following definitions.

*Stillness.* A stationary posture, with no locomotion, rearing, or grooming.

*Moving.* Translation across the chamber with a coordinated stepping gait.

*Rearing.* Extension onto the hind limbs with both forepaws off the floor, free-standing or against the chamber wall.

*Facial grooming.* Stroking of the face and ears with one or both forepaws raised to the snout.

*Body grooming.* Grooming of the torso, flank, back, or belly with the forepaws or mouth, often with the body twisted.

*Hind-paw licking.* Licking or biting of the plantar surface of the hind paw, usually while sitting.

*Scratching.* Rapid, repetitive strokes of a raised hind paw against the head, neck, or flank.

*Jumping.* A push-off from both hind limbs launching all four paws off the floor.

Each behavior was scored by an independent binary XGBoost classifier [11], with scale_pos_weight set from that behavior’s class balance. Hyperparameters were selected per behavior by Optuna [1] over 20 trials maximizing average precision. Boosting rounds were set by early stopping, 50 rounds against the held-out fold, and the final model was refit on all data at the mean best iteration across folds plus 5%. The operating threshold was chosen on the cross-validated predictions. A fixed random seed was used throughout.

Each classifier emitted a per-frame probability, which was thresholded and passed through a three-parameter filter based on BAREfoot [4], applied in order: runs shorter than min_bout were dropped, gaps up to max_gap were bridged, and a min_after_bout refractory was applied.

Settings for each classifier are given in Table S1.

### 2.6. Classifier evaluation

*Annotation dataset.* Each classifier was trained and evaluated on frames annotated frame by frame by trained observers and independently reviewed, 1,872,860 frames reviewed in total of which 167,317 were positive, over five to eight sessions per behavior. Positive-frame prevalence ranges from 1.32% to 39.4% for these behaviors. Per-behavior session counts, annotation effort, prevalence and operating points are given in Table S1. Each session was a separate mouse, and only frames a human reviewed contributed to training or evaluation.

*Evaluation scheme.* Performance was evaluated leave-one-session-out across all annotated sessions. Splitting by mouse rather than by frame avoids the data leakage that inflates reported performance when correlated frames from one mouse appear in both training and test sets [18,22]. Precision, recall, and F1 were computed per held-out session after filtering and then averaged across sessions rather than pooled across frames. Between-session spread is reported alongside the mean. To interpret each classifier, we computed SHapley Additive exPlanations (SHAP) values [25] and reported both the features with the largest mean absolute contribution and their aggregation by feature family.

### 2.7. State occupancy

For the behavioral battery, every frame was assigned to at most one state by a fixed priority order so that states cannot double-count. The order was licking, scratching, rearing, body grooming, facial grooming, moving, then stillness, giving seven mutually exclusive states. In the naloxone-precipitated withdrawal cohort, the only cohort in which jumping was scored, jumping was ranked above licking and forms an eighth state. Frames matching no classifier formed a residual unscored category. Occupancy was measured in non-overlapping 30 s bins. Panels are truncated to a fixed analysis window of 30 min, extended to 60 min for the formalin and oxycodone cohorts so that the second phase of the formalin response is included.

### 2.8. Behavioral sequencing

Using trained classifiers, we evaluated whether behavioral sequencing changed under different conditions.

*Transition tables.* Each session was reduced to its ordered list of labeled events, with consecutive identical entries collapsed and an intervening unscored stretch treated as a direct hand-off between the labeled states on either side. This gave one transition table per mouse, 7 × 7 for the seven states defined above, in which each cell counted how often one state was followed by another.

*Residuals.* Expected counts were obtained under a quasi-independence model by iterative proportional fitting to match both observed margins, with the diagonal held as a structural zero because consecutive repeats were collapsed, and each cell was expressed as a Pearson residual, (observed − expected)/√expected. Conditioning on the margins in this way removed the contribution of how often each behavior occurred, leaving first-order sequencing, meaning which behavior tended to follow which.

*Comparison between groups.* Every mouse was subsampled to the cohort’s lowest transition total, because a Pearson residual scales with the square root of the table total. Mice contributing fewer than 100 transitions were excluded: a mouse that spends most of the session in a single state, for example largely still, switches behavior rarely, and too few transitions leave its residual table too sparse to estimate. Mice were then compared by Euclidean distance between residual tables, and groups by PERMANOVA on that distance matrix, using the Behrens-Fisher variant of the pseudo-F statistic [3], with permutation restricted within animal for within-animal cohorts. The permutation spaces are small enough to enumerate exhaustively, so every reported permutation p value is exact rather than Monte Carlo sampled. For ordination, each animal’s residual table was flattened to its 42 off-diagonal cells and the animals were ordinated by principal-component analysis of those cells. Because PERMANOVA uses the Euclidean distance between the same 42-vectors, this is equivalent in Euclidean geometry to principal-coordinate analysis on that distance; the PCA form is reported because its axes have loadings, so the transitions that drive each axis can be named. Homogeneity of multivariate dispersion was checked with PERMDISP [2]. Full detail of the residual construction, the network display rules, the calibration checks and the sensitivity analyses is given in Supplementary Methods.

### 2.9. Paw contour extraction and contour metrics

Paw contours were extracted from the NIR image using pose estimation rather than from classifier output. Within a 100 × 100 pixel region of interest centered on each hind-paw keypoint, the grayscale frame was smoothed with a 3 × 3 Gaussian kernel and binarized by Otsu’s method [30], computed independently for each region of interest on each frame, and the largest external contour taken as the paw. Each metric was expressed as the ratio of the injured or injected hind paw (HL) to the contralateral hind paw (HR).

*Contour intensity* is the mean grayscale value of the pixels enclosed by the contour. Because the NIR light reflected back to the camera increases with the pressure a paw exerts against the acrylic, intensity provides an optical index related to paw contact rather than a calibrated measure of load.

*Contour area* is the area enclosed by the contour and reflects the projected size of the contact footprint. A paw that is lifted, curled, or bearing less weight presents a smaller contacting surface.

*Contour circularity* is defined as 4πA/P², where A is the contour area and P its perimeter. It equals 1 for a perfect circle and decreases as the outline becomes elongated or its perimeter irregular.

*Contour solidity* is defined as A/A_hull, where A_hull is the area of the convex hull of the contour. It equals 1 for a fully convex outline and falls below 1 in proportion to the concavities present, such as the notches between extended toes.

Contours and frames were filtered before contributing to the index, including exclusion of frames scored as licking, during which the paw leaves the floor and the contact signal is lost. The filters and the frame retention they produce are given in Supplementary Methods.

### 2.10. Statistics

Two independent groups were compared by Welch’s unequal-variance t test, within-animal comparisons by paired t test, dose-response experiments by one-way ANOVA with Welch t tests against the common vehicle control and Holm correction within each dose family, binned time courses by mixed-design repeated-measures ANOVA, with group as the between-animal factor and 5 min bin as the within-animal factor and Greenhouse-Geisser correction applied to the within-animal terms, monotonic dose trends by Spearman correlation, apparatus comparisons by Mann-Whitney, and sequencing by PERMANOVA. Dose-response curves were normalized to the vehicle mean within each phase and fit by hyperbolic regression with the maximum constrained to 100% to derive the ED_50_ with a 95% confidence interval, as described for hand-scored data in this assay [31]. Effect sizes are reported as Hedges g for unpaired comparisons, Cohen dz for paired comparisons, and η² or partial η² for ANOVA terms. Analyses were performed in Python 3.10.6 with numpy 1.26.4, scipy 1.15.3, statsmodels 0.14.2, pandas 2.2.3, scikit-learn 1.4.2, xgboost 2.1.4, opencv 4.8.1 and matplotlib 3.10.9, and pose estimation used DeepLabCut 3.0.0rc13. All analyses were run on a Windows 11 workstation (Intel Core i9-12900K, 64 GB RAM, NVIDIA GeForce RTX 3080 Ti with 12 GB of video memory); PixelPaws runs on a standard Windows PC, but an NVIDIA GPU is recommended because pose estimation runs substantially slower on the CPU alone. Data are expressed as mean +/- SEM with individual animals overlaid, significance was set at p < 0.05, and exact statistics and post hoc comparisons are given in the table accompanying each figure.

### 2.11. Data and code availability

PixelPaws, including the analysis interface and the apparatus design files and STLs, is released under a CC BY-NC 4.0 license at https://github.com/rslivicki/PixelPaws. The bundled pretrained pose network and the reference behavior classifiers are distributed as tagged release assets.

PawCapture, the multi-camera acquisition application, is distributed in the same repository under pawcapture/ as a bundled Windows executable that requires no separate Python installation.

The analysis interface also exposes the frame-labeling and network-training steps used to produce the bundled pose network, so a laboratory whose recordings differ from the reference condition, for example in coat color, can add labeled frames and train a new network through the same interface.

Every figure is produced by a script deposited with the code. Analysis caches are written alongside the figures, so that a rebuild re-plots without rescoring. Raw video and per-frame prediction files are available from the corresponding author on reasonable request.

## 3. Results

### 3.1. The PixelPaws platform resolves per-frame pose, paw brightness and behavior

Mice were recorded from a bottom-up view through a transparent acrylic floor under infrared illumination, in parallel chambers controlled from a single laptop (Fig. 1A). Acquisition was handled by PawCapture, custom multi-camera software that records synchronized video with calibrated spatial scaling (Fig. 1B), and all downstream processing ran in PixelPaws, an integrated analysis application covering pose estimation, feature extraction, classifier training, prediction review and gait analysis (Fig. 1C). For each frame, markerless pose estimation predicted nine body parts, and paw brightness was extracted from the region surrounding each paw keypoint (Fig. 1D). Pose and brightness were converted into a per-frame feature matrix comprising joint angles, inter-part distances, velocities and paw brightness (Fig. 1E), which a supervised classifier converted into a per-frame probability for each behavior. Contiguous frames whose probability exceeded that classifier’s operating threshold were merged into events (Fig. 1F). Using this approach, we generated eight supervised classifiers: stillness, moving, rearing, facial grooming, body grooming, hind-paw licking, scratching and jumping.

Hind-paw licking, scratching, and jumping were used for the pharmacological experiments below and are characterized in the main figures. Attribution for the remaining five was consistent with the physical definition of each behavior. Feature-family attribution for all five is shown in Figure S1A,D,G,J,M, with SHAP values for the strongest individual features (Fig. S1B,E,H,K,N) and performance across decision thresholds (Fig. S1C,F,I,L,O) alongside (Table S1).

### 3.2. Classifier output is comparable across two recording setups

During development, the current-limiting resistors on the LED strip were found to warm the acrylic floor, which could itself alter behavior. Lowering the LED supply voltage from 12 V to 5 V removed the resistive warming of the floor, with a compensating change to exposure settings. To ask whether this affected the behavioral readout, ten mice were recorded, six on the first setup, which used the voltage step-down, and four on the second, which did not, each contributing one recording in one setup, and percent time in stillness, moving, rearing, facial grooming, body grooming, licking and scratching was compared between setups. No behavior differed between setups (p = 0.35 to 1.00, Mann-Whitney; Fig. S2A-G; Table S2), with differences of 0.34 to 2.85 percentage points of the session, and the stillness time course was comparable across recordings (Fig. S2H).

### 3.3. Classifier and contour readouts survive video compression

Raw video is large and scales poorly with cohort size, so we asked whether we could compress video files without loss of pose-estimation and paw contour accuracy. A 3-min segment (minutes 5 to 8) of one session from each of four mice was re-encoded from the source recording, the hardware-encoded H.264 file written at acquisition (∼8 Mbit/s) at five x264 rates and one x265 rate, and each encode was run through the full pipeline independently of the others. Compression level was set by the constant rate factor (CRF), on which lower values mean less compression and a larger file. Deviation from the source recording is shown for bitrate, pose keypoint position, paw contour brightness and the contour metrics introduced below, together with frame-by-frame agreement of the behavioral state call and the change in whole-session behavioral composition (Fig. S3A-F; Table S3). At CRF 23, the setting used for all analyses in this manuscript, the bitrate was 23 times lower than the source (0.35 against 8.1 Mbit/s) while pose moved 0.48 px, paw brightness changed 0.28%, the contour intensity ratio shifted 0.001 log units, and the behavioral state call agreed with the source recording on 91.8% of frames. Thus, compression at this rate had negligible effects on tracking and contour endpoints and left classifier output largely unchanged.

### 3.4. A supervised classifier detects hind-paw licking and reports increased licking after formalin

We trained a classifier to detect hind-paw licking, a principal nocifensive response in inflammatory pain models and conventionally scored in the formalin assay [15,38] (Fig. 2A). Feature-family attribution indicated that body angle contributed most to classification, followed by inter-part distance and velocity (Fig. 2B), with SHAP values for individual features shown by beeswarm (Fig. 2C). The classifier reached an out-of-fold F1 of 0.84 (precision 0.83, recall 0.85) by leave-one-session-out evaluation, with performance across decision thresholds and agreement with human annotation shown in Figure 2D,E (Table S1).

**Figure 2.**
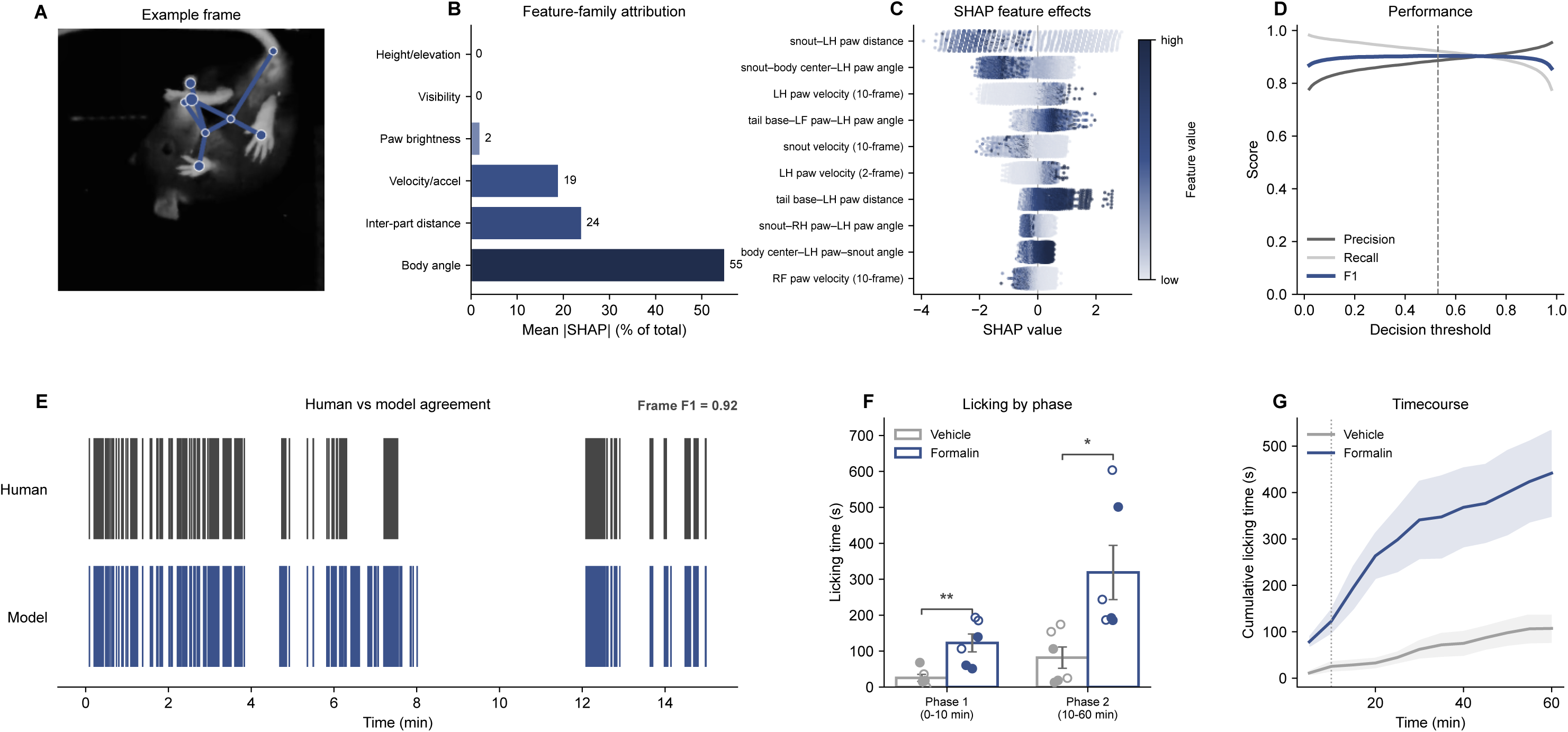
A supervised classifier detects hind-paw licking and reports increased licking after formalin. A representative licking frame is shown (A). Feature-family attribution indicated that body angle contributed most to classification, followed by inter-part distance and velocity, with paw brightness contributing little to this behavior (B), and SHAP values for the individual features are shown by beeswarm (C). Classifier performance across decision thresholds is summarized by precision, recall and F1 (D), and agreement with human annotation is shown as an ethogram over a held-out session, with the frame-level F1 for that session (E). The operating threshold was chosen on cross-validated predictions, not on the held-out session. When the classifier was applied to formalin- and vehicle-injected animals, formalin increased licking in both phases of the response (F), with the expected biphasic time course (G). ****p<0.0001, ***p<0.001, **p<0.01, *p<0.05 Welch t test (F). Post-tests and other statistics are reported in the per-figure statistics table. Data are represented as mean +/- SEM. N = 6 per group.

We next evaluated whether this trained classifier detected a pharmacologically evoked change in licking. Mice received an intraplantar injection of formalin or saline vehicle into the left hind paw and were recorded for 60 min. Formalin injection increased total licking time relative to vehicle (442 ± 92.5 s versus 107 ± 29.7 s, t_6.0_ = 3.443, p = 0.0137, Hedges g = 1.99, Welch t; Table 2), and the increase was present in both the acute phase 1 (0 to 10 min) and the prolonged inflammatory phase 2 (10 to 60 min) (Fig. 2F; Table 2). Cumulative licking was higher in formalin-injected mice throughout the session (group F_1,10_ = 12.908, p = 0.0049, partial η² = 0.563; group by bin interaction F_11,110_ = 6.129, Greenhouse-Geisser corrected p = 0.0198; mixed-design repeated-measures ANOVA; Fig. 2G; Table 2). Rescoring the same recordings with each behavioral classifier indicated that this increase occurred against an otherwise largely unchanged behavioral background: licking occupied 2.39% of the session in vehicle-injected mice and 12.37% after formalin, while the remaining behaviors shifted comparatively little (Fig. S4). The classifier therefore recovers a temporal signature established by manual scoring, and the change is specific to licking rather than reflecting a general shift in behavior.

### 3.5. Formalin reduces hind-paw contour intensity and alters the shape of the injected paw

We next asked whether the same recordings could capture paw-level changes accompanying formalin-evoked behavior. Hind-paw contours were extracted from the formalin cohort in Figure 2 and used to quantify intensity, area, circularity, and solidity (Fig. 3A). Each metric was expressed as the ratio of the injected to the contralateral paw, such that a value of 1 indicates symmetry between the two paws. The contour intensity ratio fell below 1 in formalin-injected mice and remained depressed across the session, whereas vehicle-injected mice remained near 1 (group F_1,10_ = 17.211, p = 0.0020, partial η² = 0.633, mixed-design repeated-measures ANOVA; Fig. 3B; Table 3). The size of the difference varied across the session (group by bin interaction F_11,110_ = 2.730, Greenhouse-Geisser corrected p = 0.0407), so the four contour metrics (intensity, area, circularity and solidity) were compared by phase. The injected paw was dimmer than the contralateral paw in both phases, at 0.902 against 1.00 in phase 1 and 0.915 against 1.01 in phase 2. The injected paw was also rounder and more solid during phase 1 only, while area was unchanged throughout (Welch t; Fig. 3C,D; Table 3). The intensity difference was present in both phases while the shape change was confined to the acute phase. The contour therefore yields paw-level metrics alongside the behavior-level labels.

**Figure 3.**
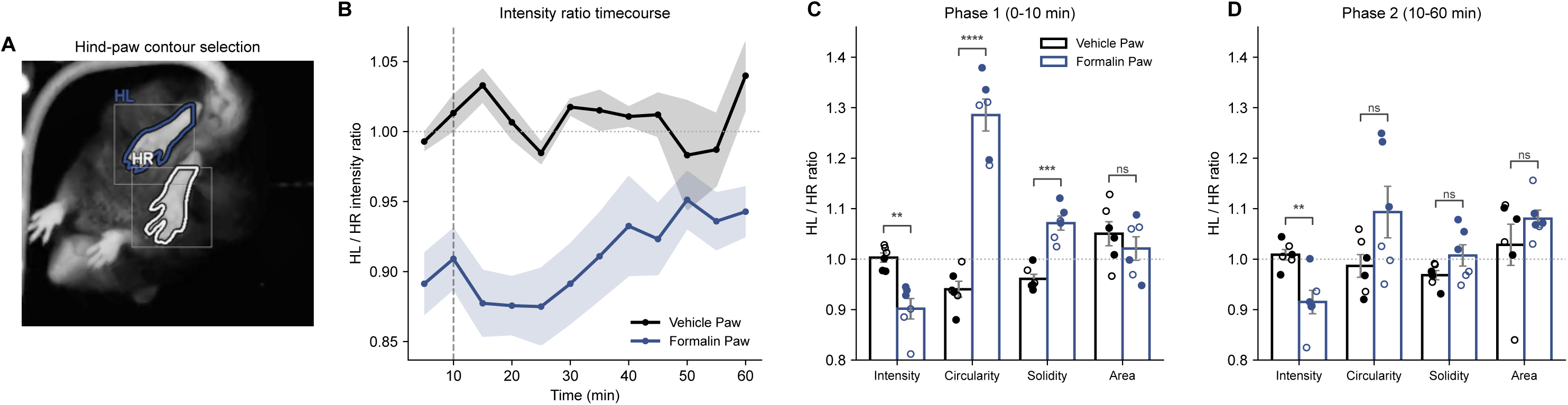
Formalin reduces hind-paw contour intensity and alters the shape of the injected paw. Hind-paw contours were extracted within a region of interest centered on each hind-paw keypoint and expressed as ratios of the injected (HL) to contralateral (HR) paw, so that a value of 1 indicates symmetry. A representative contour selection is shown (A). The HL/HR contour intensity ratio fell below 1 in formalin-injected animals and remained depressed across the session, whereas vehicle-injected animals remained near 1 (B). The injected paw was dimmer in both phases, and rounder and more solid in phase 1 only; contour area was unchanged (C, D). ****p<0.0001, ***p<0.001, **p<0.01, *p<0.05 Welch t test (C, D). Post-tests and other statistics are reported in the per-figure statistics table. Data are represented as mean +/- SEM with individual animals overlaid. N = 6 per group.

### 3.6. Oxycodone dose-dependently reduces formalin-evoked licking and paw contour asymmetry

We next used the licking classifier to construct a dose-response for oxycodone, a known analgesic in the formalin assay [31]. Mice received oxycodone (1, 3 or 10 mg/kg s.c.) or vehicle before intraplantar formalin and were recorded for 60 min. Cumulative licking is shown over the full session (Fig. 4A). Oxycodone reduced licking dose-dependently across the whole session (F_3,19_ = 12.475, p < 0.0001), in phase 1 (F_3,19_ = 12.444, p < 0.0001) and in phase 2 (F_3,19_ = 9.758, p = 0.0004; one-way ANOVA effect of dose, with Welch t against vehicle as post-test; Fig. 4B,C; Table 4). Whole-session licking was reduced at 10 mg/kg but not at 3 or 1 mg/kg (Table 4).

**Figure 4.**
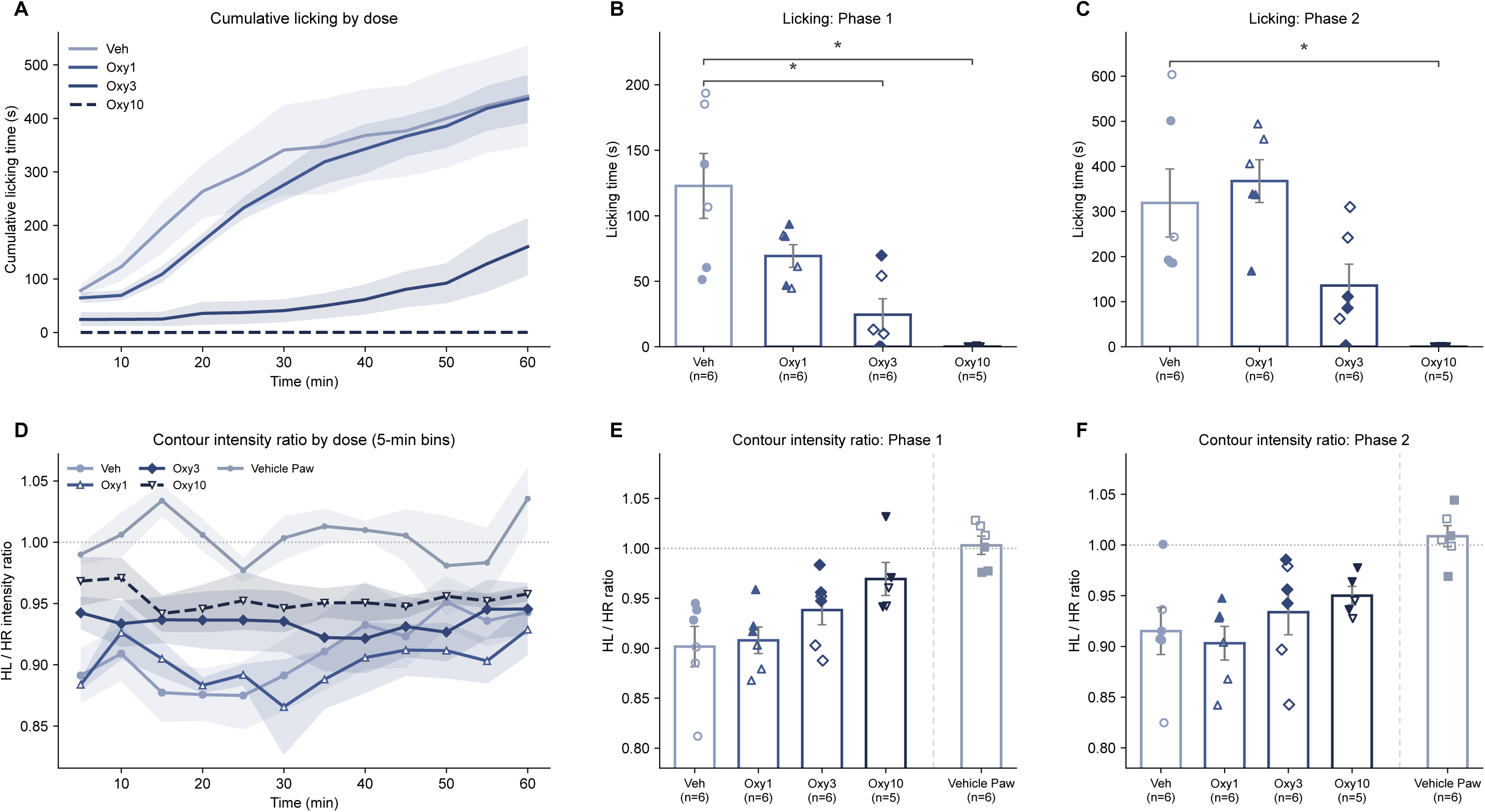
Oxycodone dose-dependently reduces formalin-evoked licking and paw contour asymmetry. Animals received oxycodone (1, 3, 10 mg/kg s.c.) or vehicle before formalin, and licking and paw contour intensity were measured across the session. Cumulative licking is shown over the full session (A) and summarized for phase 1 (0-10 min; B) and phase 2 (10-60 min; C). The HL/HR contour intensity ratio is shown as a five-minute time course (D) and summarized for phase 1 (E) and phase 2 (F). ****p<0.0001, ***p<0.001, **p<0.01, *p<0.05 Welch t test versus vehicle, Holm-corrected within each phase (B, C, E, F); the one-way ANOVA effect of dose is reported in the per-figure statistics table with the other statistics. The state traces (B,E,H,K) show mean +/- SEM; rasters and cumulative curves show individual animals, with the group mean overlaid on the cumulative panels. N = 5-6 per group.

We next evaluated broad behavioral changes across all of our classifiers. Time spent still fell across the dose series, from 30.4% of the session in vehicle-treated mice to 0.2% at 10 mg/kg, while time spent moving rose from 18.2% to 96.7% (Fig. S5), the direction expected of a mu-opioid agonist [35].

Manual scoring of this assay has established a potency benchmark for oxycodone [31]. Phase 2 estimates were comparable, 3.27 mg/kg against the reported 3.4 mg/kg, and restricting phase 2 to the 16 to 40 min window reported in that paper gave 3.34 mg/kg. The whole-session estimate was 2.52 mg/kg. Our phase 1 estimate was lower than reported, 0.99 against 2.7 mg/kg. That study gave drugs i.p. 45 min before 5% formalin, whereas here oxycodone was given s.c. 5 min before 2% formalin, and the differences in pretreatment interval, route and formalin concentration may account for the lower phase 1 estimate. Confidence intervals for the phase 1 and phase 2 fits are given in Table S4. The classifier therefore recovers a phase 2 potency estimate for oxycodone comparable to that obtained by blinded manual scoring of the same assay.

Contour intensity tracked the dose series in phase 1. Across doses, the injected paw to contralateral paw intensity ratio moved toward symmetry with increasing dose (rho = +0.58, p = 0.0034, Spearman), an effect confined to the acute phase and absent in phase 2 (rho = +0.35, p = 0.1060). The index is shown as a five-minute binned time course (Fig. 4D) and summarized by phase (Fig. 4E,F; Table 4), with a vehicle paw-injection reference group in panels 4E and 4F. Contour intensity therefore tracks the pharmacological response across the dose range rather than only the presence of injury.

### 3.7. Oxycodone reduces licking and shifts hind-paw contour area toward symmetry after spared nerve injury

We next asked whether oxycodone would suppress spontaneous licking and paw contour asymmetry in a mouse model of chronic neuropathic pain. Male mice that had undergone spared nerve injury were recorded at baseline and again following oxycodone (3 mg/kg s.c.) or vehicle, and the first 30 min post-drug were analyzed. Cumulative licking was lower in oxycodone-treated mice (19.9 ± 6.8 s versus 106 ± 17.6 s, t_7.8_ = -4.575, p = 0.0019, Hedges g = -2.45, Welch t; Fig. 5A; Table 5).

**Figure 5.**
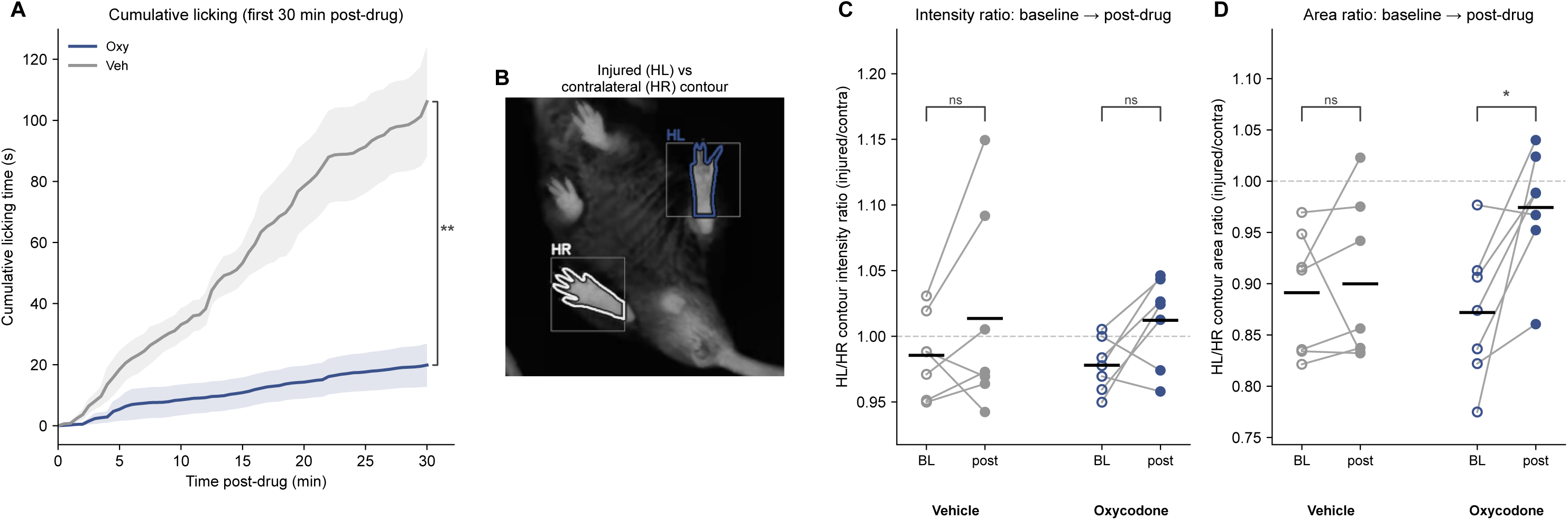
Oxycodone reduces licking and shifts hind-paw contour area toward symmetry after spared nerve injury. Spared nerve injury animals were recorded at baseline and again following oxycodone (3 mg/kg s.c.) or vehicle, and the first 30 min post-drug were analyzed. Cumulative licking was lower in oxycodone-treated animals (A). A representative frame is shown with the injured (HL) and contralateral (HR) hind-paw contours drawn on it (B). Because each animal serves as its own control, the HL/HR intensity ratio (C) and area ratio (D) are shown as paired baseline-to-post-drug comparisons, with open symbols denoting baseline and filled symbols post-drug. ****p<0.0001, ***p<0.001, **p<0.01, *p<0.05 paired t test (C,D); Welch t test (A). Post-tests and other statistics are reported in the per-figure statistics table. The time course (A) shows mean +/- SEM; C and D show individual animals with group means. N = 7 per group, all male.

Oxycodone shifted the area ratio toward symmetry, from 0.872 ± 0.025 to 0.974 ± 0.022 (t_6_ = 3.341, p = 0.0156, Cohen dz = 1.26, paired t; Fig. 5B,D), with the intensity ratio trending in the same direction (0.978 ± 0.008 to 1.012 ± 0.013; t_6_ = 2.345, p = 0.0575; Fig. 5C). Neither ratio changed in vehicle-treated mice (area p = 0.7403; intensity p = 0.2260; Fig. 5C,D; Table 5). The change from baseline differed between treatments for the area ratio (oxycodone +0.102 ± 0.031 versus vehicle +0.009 ± 0.025, t_11.5_ = 2.381, p = 0.0356, Hedges g = 1.27, Welch t) but not for the intensity ratio (t_10.8_ = 0.245, p = 0.81; Table 5). As in the formalin dose-response, oxycodone raised locomotion rather than producing general behavioral suppression, with time moving rising from 30.1% at baseline to 76.9% post-drug while vehicle-treated mice fell from 37.0% to 20.9% (Fig. S6H,K).

### 3.8. A supervised classifier detects scratching

We next trained a second classifier to detect hindlimb scratching, a common readout in pruritogenic assays (Fig. 6A). Feature-family attribution indicated that velocity and acceleration contributed most to this classifier, ahead of body angle and paw brightness (Fig. 6B), with SHAP values for individual features shown by beeswarm (Fig. 6C). The classifier reached an out-of-fold F1 of 0.75 (precision 0.84, recall 0.70), with performance across decision thresholds and agreement with human annotation shown in Figure 6D,E (Table S1).

**Figure 6.**
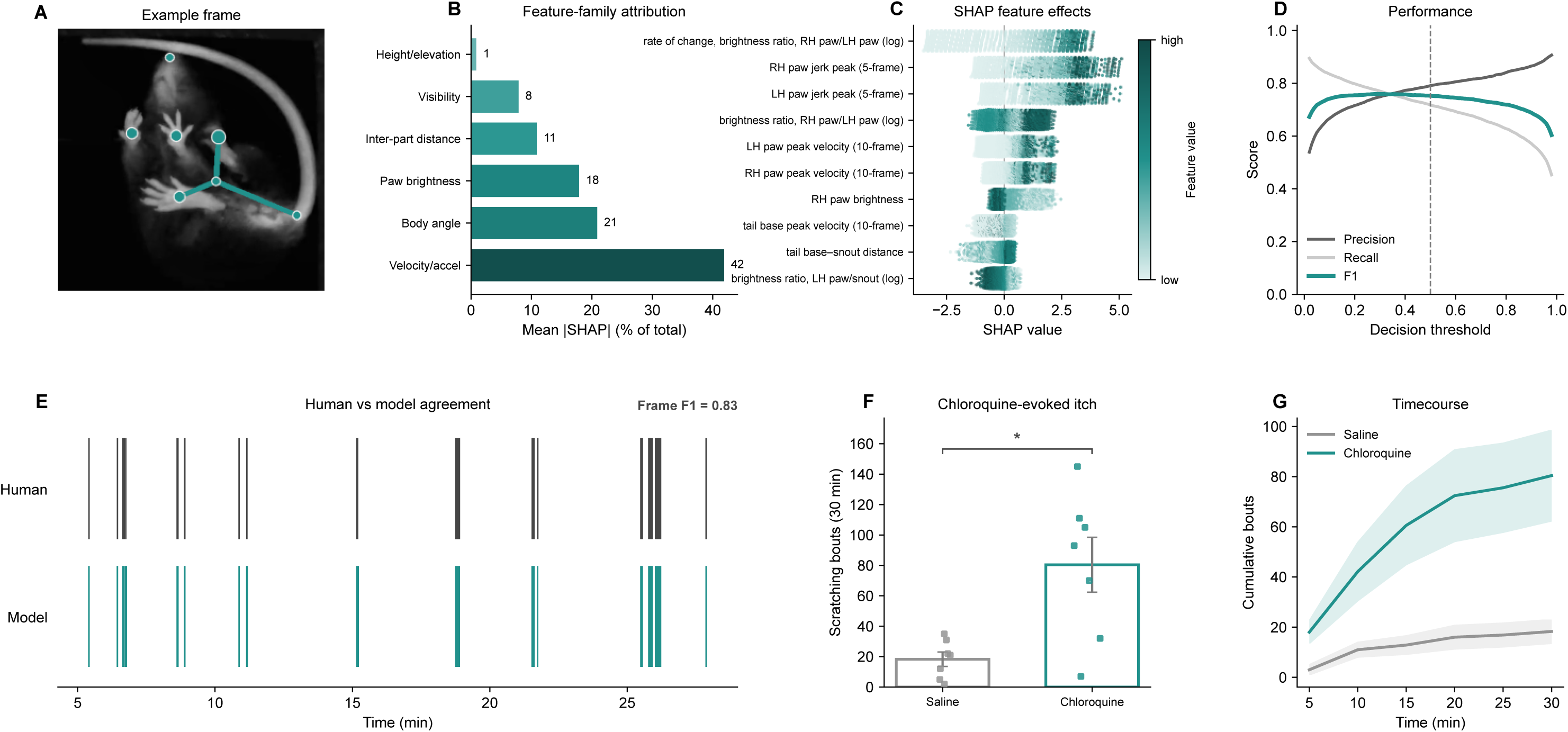
A supervised classifier detects scratching. A representative scratching frame is shown (A). Feature-family attribution is shown for the scratching classifier (B) with SHAP values for individual features by beeswarm (C). Classifier performance across decision thresholds is summarized by precision, recall and F1 (D), and agreement with human annotation is shown as an ethogram over a held-out session, with the frame-level F1 for that session (E). The classifier was then applied without retraining to an independent cohort receiving chloroquine or saline, analyzed over the first 30 min post-injection at the shipped operating point (probability 0.5, minimum bout 3 frames, maximum gap 10 frames): chloroquine-treated animals showed more scratching (F), which accumulated more rapidly across the session (G). ****p<0.0001, ***p<0.001, **p<0.01, *p<0.05 Welch t test (F). Post-tests and other statistics are reported in the per-figure statistics table. Data are represented as mean +/- SEM with individual animals overlaid. N = 7 per group, all female (F,G).

The scratching classifier was then used to evaluate chloroquine-induced itch, with each mouse recorded at baseline and again after injection of chloroquine or saline. Chloroquine increased scratching over the 30 min following injection, from 0.235 ± 0.088% of the session at baseline to 0.794 ± 0.197% after treatment (t_7_ = 2.610, p = 0.0349, Cohen dz = 0.92, paired t), whereas saline-treated mice did not change (Table 6). The change from baseline differed between groups (t_8.2_ = 2.701, p = 0.0266, Hedges g = 1.31, Welch t), and both scratching bouts (80.4 ± 18.1 versus 18.3 ± 4.7, t_6.8_ = 3.327, p = 0.0131; Fig. 6F) and scratching time (39.6 ± 11.4 s versus 10.3 ± 3.0 s, t_6.8_ = 2.487, p = 0.0427; Table 6) over the window were higher after chloroquine. Cumulative scratching bouts were higher throughout the 30-min window (treatment F_1,12_ = 10.559, p = 0.0070, partial η² = 0.468; treatment by bin interaction F_5,60_ = 5.619, Greenhouse-Geisser corrected p = 0.0221; mixed-design repeated-measures ANOVA; Fig. 6G; Table 6), the interaction reflecting a difference that opened within the first 10 min and then widened more slowly. Scoring the full repertoire showed no major differences between chloroquine- and saline-treated mice in any other behavior (Fig. S7).

### 3.9. Rimonabant evokes dose-dependent scratching

We next tested the scratching classifier against a second pruritogen acting through a different mechanism. Rimonabant, a CB1 receptor antagonist, evokes dose-dependent scratching independently of histamine [13,34]. Scratching was scored in mice receiving rimonabant (1, 3 or 10 mg/kg, i.p.) or vehicle over 60 min. Total scratching increased dose-dependently (F_3,23_ = 5.825, p = 0.0041, one-way ANOVA effect of dose, with Welch t against vehicle as post-test; Fig. S8A; Table S5), with the corresponding cumulative time course (Fig. S8B), and the increase occurred against an otherwise stable repertoire (Fig. S9).

### 3.10. A supervised classifier detects jumping and reports naloxone-precipitated opioid withdrawal

Jumping is among the most frequently scored somatic signs of opioid withdrawal in mice [8,9,12]. Using a model of acute oxycodone-induced withdrawal, in which mice received a single dose of oxycodone (9 mg/kg s.c.) and withdrawal was precipitated 2 h later with naloxone (3 mg/kg s.c.), we trained a jumping classifier. Paired frames show a mouse immediately before and during a jump (Fig. 7A). Feature-family attribution indicated that paw brightness and velocity contributed most to classification (Fig. 7B), consistent with loss of floor contact during the airborne phase, with SHAP values for individual features shown by beeswarm (Fig. 7C). The classifier reached an out-of-fold F1 of 0.92 (precision 0.95, recall 0.89), with performance across decision thresholds and agreement with human annotation shown in Figure 7D,E (Table S1).

**Figure 7.**
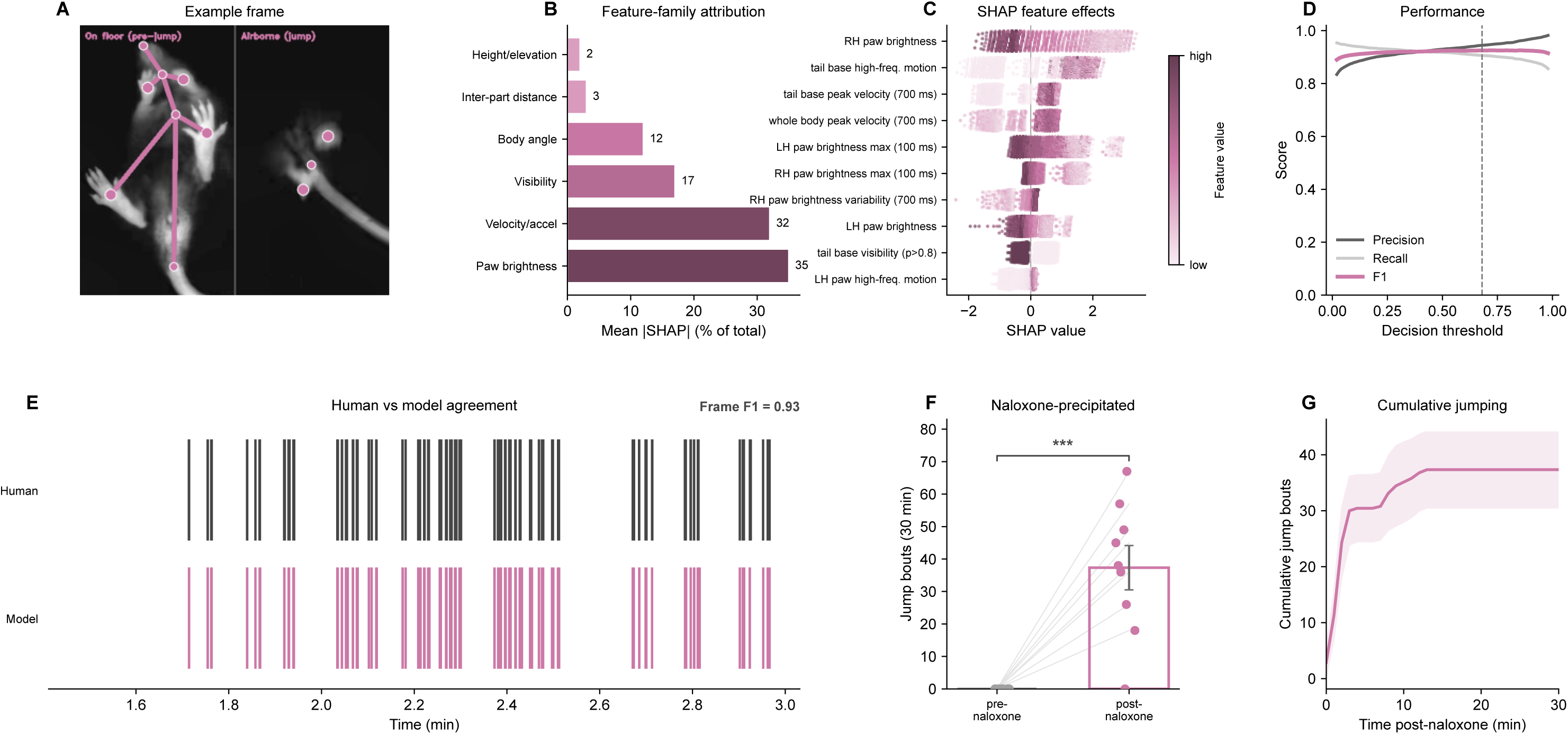
A supervised classifier detects jumping and reports naloxone-precipitated opioid withdrawal. Paired frames show a mouse immediately before and during a jump (A). Feature-family attribution indicated that paw brightness and velocity contributed most to classification (B), consistent with loss of contact with the acrylic during the airborne phase, with SHAP values for individual features by beeswarm (C). Classifier performance across decision thresholds is summarized by precision, recall and F1 (D), and agreement with human annotation is shown as an ethogram over a held-out session, with the frame-level F1 for that session (E). In oxycodone-treated animals, naloxone precipitated a marked increase in jumping relative to the pre-naloxone period (F), concentrated in an acute burst following injection (G). ****p<0.0001, ***p<0.001, **p<0.01, *p<0.05 paired t test (F). Post-tests and other statistics are reported in the per-figure statistics table. The time course (G) shows mean +/- SEM; F shows individual animals with group means. N = 9, all female.

Naloxone precipitated an increase in jumping relative to the pre-naloxone period, from none before injection to 37.3 ± 6.8 events after (t_8_ = 5.465, p = 0.0006, Cohen dz = 1.82, paired t; Fig. 7F; Table 7), concentrated in an acute burst following injection (Fig. 7G; Table 7). Scoring the whole repertoire indicated the mirror image of the oxycodone dose series: time moving fell from 87.9% to 15.7% and time still rose from 1.6% to 41.6% after naloxone, with jumping occupying 1.0% of the post-naloxone session against that background (Fig. S10).

### 3.11. Naloxone-precipitated withdrawal alters behavioral organization

We next asked whether withdrawal alters the order in which behaviors follow one another, independently of how much of each occurs. The Figure 7 recordings were rescored with the seven classifiers that partition the session into mutually exclusive states and read as sequence rather than occupancy. Jumping, the eighth classifier and the readout of Figure 7, is brief and overlaps the other states, so it was analyzed as events there rather than as a state here.

Withdrawal reorganized the transition structure. Behaviors are drawn as a network before naloxone (Fig. 8A) and during withdrawal (Fig. 8B), with edge color giving the Pearson residual against each group’s own margins, so that a behavior becoming more common cannot move an edge. Eight of 20 assessable routes changed by more than four Pearson-residual units (Fig. 8C). Body grooming is drawn dashed because it supplies too few exits to be assessable (Supplementary Methods). The two conditions separated in ordination (pseudo-F = 7.455, R² = 0.318, p = 0.0039, the smallest value attainable for nine paired animals, PERMANOVA restricted within animal, against a 95th percentile of the permutation R² distribution of 0.18 at this group size; Fig. 8D; Table 8).

**Figure 8.**
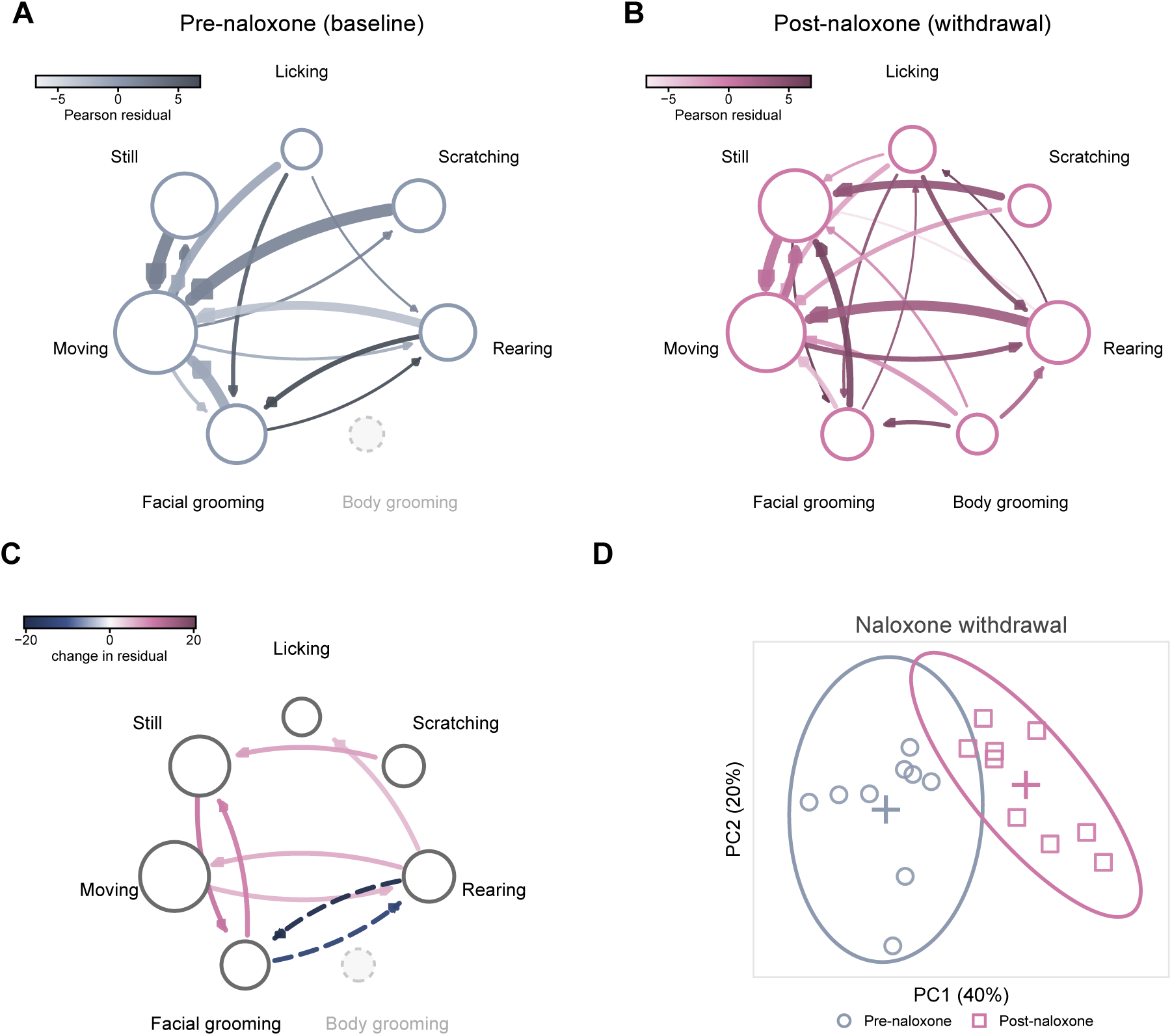
Naloxone-precipitated withdrawal alters behavioral organization. Recordings from Figure 7 were rescored with the seven classifiers that partition the session into mutually exclusive states (licking, scratching, rearing, body grooming, facial grooming, moving and stillness) and read as sequence. Behaviors are drawn as networks before naloxone (A) and during withdrawal (B); node size indicates time spent, edge width the share of that state’s exits, and edge color the Pearson residual from a quasi-independence fit. Routes that changed between conditions are shown in (C), solid where strengthened and dashed where weakened; body grooming is drawn dashed and unlabeled where it falls below the bout floor required to characterize a state. Each animal is placed by how differently it strings behaviors together (D); the surrounding line is a 95% normal-theory ellipse describing the spread of each group, not a confidence region for its mean. The corresponding contrast for every other cohort is in Figure S11. PERMANOVA with permutation restricted within animal (D); panels A-C carry no test. Post-tests and other statistics are reported in the per-figure statistics table. Data are represented as individual animals, open circles before naloxone and open squares during withdrawal, with group centroids as crosses (D). N = 9 female mice, each contributing both phases.

Precipitated withdrawal therefore alters the order in which behaviors follow one another, independently of how much of each occurs.

### 3.12. Behavioral sequencing distinguishes the state produced by each manipulation

We next applied the same test to every contrast in the battery. Formalin altered behavioral sequencing (pseudo-F = 3.019, R² = 0.232, p = 0.0281). While oxycodone diminished formalin-induced licking (Fig. 4B,C), the remaining repertoire collapsed onto locomotion rather than resembling that of vehicle paw-injected mice, with every state exiting into moving, time spent moving rising from 18.2% of the session in vehicle-treated mice to 96.7% at 10 mg/kg, and stillness falling from 30.4% to 0.2% (Fig. S5). Placing the oxycodone dose series in the same space as a vehicle paw-injection reference, distance from the vehicle-paw centroid rose with dose rather than falling (rho = +0.52, p = 0.0122; Fig. S11B), so higher doses moved sequencing further from the vehicle pattern rather than back toward it. The same collapse appeared in oxycodone-treated nerve-injured mice (pseudo-F = 25.579, R² = 0.681, p = 0.0006). Oxycodone-treated mice were also half as dispersed as vehicle-treated mice (mean distance to group centroid 2.19 versus 4.26, PERMDISP p = 0.0006; Table 8), consistent with every animal converging on the same locomotor pattern. No other two-group contrast differed in dispersion (PERMDISP p = 0.075 to 0.79), although dispersion differed across the five groups of the dose series (F = 5.185, p = 0.018), with the 10 mg/kg group the tightest (mean distance to centroid 3.68 against 5.03 to 7.64; Table 8). Chloroquine raised scratching roughly threefold over the same window but left the transitions between behaviors intact (pseudo-F = 0.512, p = 0.8594), whereas rimonabant produced a trend toward reorganization that did not reach significance (pseudo-F = 2.029, R² = 0.169, p = 0.0563, against a 95th percentile of the permutation R² distribution of 0.17; Fig. S11A,C-F; Table 8).

These data indicate that suppression of a target behavior can occur without restoration of behavioral organization.

## 4. Discussion

Here we present an integrated platform for automated scoring of mouse behavior, combining a low-cost hardware apparatus, video acquisition software, and a graphical analysis application shipped with a pretrained pose-estimation network and eight behavioral classifiers. Applied across noxious, pruritic, and drug-dependent states, classifiers for licking, scratching, and jumping agreed with trained human observers and recovered a published oxycodone potency estimate from the formalin assay. The same recordings also yielded continuous paw contour metrics and measures of behavioral sequencing.

Many tools now exist for generating behavioral classifiers and have substantially lowered the barrier to automated behavior analysis [7,18,19,21]. Adopting them nonetheless carries a substantial learning curve: installing and reconciling the Python dependencies for pose estimation and for the classifier packages is a recurring obstacle, and published hardware builds add further financial and time commitments. Standardizing the apparatus is intended to make recordings comparable across builds, so the models can be bundled pretrained and the platform used as an out-of-the-box scoring pipeline. The apparatus described here requires three 3D-printed parts (chamber, lid and camera mount), an LED strip light, a near-infrared camera and lens, and made-to-order acrylic panels, for under $300 per enclosure in materials, and the printed parts can be ordered from commercial printing services where a 3D printer is not available. PixelPaws installs from a single package that resolves the Python dependencies without user intervention. The bundled models are specific to the reference apparatus, but the software is not: a laboratory can train its own behavioral classifiers on its own recordings and hardware through the same interface.

Feature attribution recovered the physical description of each behavior. Licking requires the mouse to bring the hind paw to the mouth, and was dominated by postural geometry, with body angle contributing most. Scratching is conventionally defined as rapid, repetitive movement of the hindlimb toward the affected site [23], and velocity and acceleration accounted for the largest share of feature attribution, ahead of body angle. Brightness and velocity dominated jumping, consistent with loss of floor contact during the airborne phase. The scratching classifier detected chloroquine-evoked scratching in one cohort and a rimonabant dose response in another, suggesting that it captures the scratching behavior itself rather than being specific to the compound used to evoke it. The same feature ordering appears in BAREfoot, where spatial features carried the highest importance for licking and biting [4].

Pharmacology studies incorporating both sexes, a range of doses, and multiple compounds generate large datasets quickly, and the resulting volume of video is impractical to score manually. A recent comparison of morphine, oxycodone and a biased mu-opioid agonist across four pain models used 435 mice, and formalin-evoked licking was scored manually in 5 min bins by observers blinded to treatment [31]. We took that design as the test case and asked whether the platform could recapitulate an oxycodone dose-response in the formalin assay. Oxycodone (1, 3 or 10 mg/kg s.c.) reduced formalin-evoked licking dose-dependently across the session, and the classifier recovered an ED_50_ comparable to that obtained by blinded manual scoring of the same assay. Here, the equivalent series was scored automatically, requiring only the operator time needed to set up the analysis, and returned a comparable phase 2 potency estimate. Other groups have used frustrated total internal reflection, in which light guided within a glass floor escapes and scatters where the paw makes contact, to infer floor contact and detect mechanical hypersensitivity [36,43], which may be informative alongside the scoring of expressed behaviors. We therefore evaluated the contour metrics in these assays, segmenting the paw contour by Otsu’s method, which yields intensity and area specific to the paw, and also returns its shape. Formalin reduced the contour intensity ratio across the session, and in nerve-injured mice oxycodone shifted the contact area ratio toward symmetry within animal, with intensity trending in the same direction. Paw luminance is also offered commercially as a weight-bearing readout [6], and the implementation described here is open and obtained from a single camera without dedicated contact optics, using the same recordings that produce the behavioral scores. The contour signal also tracked the oxycodone dose-response in phase 1, so the readout is pharmacologically sensitive as well as reporting the presence of injury.

Another strength of the approach is that multiple classifiers report both the time spent in each behavior and the transitions between them, an area of growing interest among behavioral neuroscientists, and machine-learning output has been read as sequence in other behavioral contexts [41]. Reading our classifier output the same way, the largest changes in transition structure were observed in naloxone-precipitated withdrawal and in oxycodone-treated nerve-injured mice, with a smaller change after formalin and none after chloroquine. Chloroquine raised scratching approximately threefold over the same window, so the analysis responds to reorganization of the repertoire rather than to an increase in a single behavior. However, oxycodone did not return formalin-treated mice to the vehicle pattern, and consistent with reports using LUPE [29] it increased locomotion, suppressing formalin- and injury-evoked licking while reorganizing the remainder of the repertoire around locomotion, and producing a transition pattern distinct from both formalin and vehicle. Consistent with this, English et al. [16] predicted the dose of THC a mouse had received from pose-tracked behavior, and the prediction improved as more of the behavioral profile was included. Whether a candidate analgesic normalizes behavioral organization, rather than only suppressing a target behavior, may therefore offer insight that simply scoring one behavior alone cannot.

While the software itself is not restricted to a particular setup, the bundled pose network and classifiers were developed for the standardized reference apparatus. We have not quantified agreement between sites, and the models have not been tested on other hardware. Related to this, although the classifiers were trained on more than 1.8 million reviewed frames, frames within a session are highly correlated, so the effective sample size is the five to eight annotated sessions per behavior rather than the frame count. Leave-one-session-out evaluation accounts for this, but broader validation across animals, experimenters, laboratories and recording conditions remains necessary before the bundled models can be treated as generalizing beyond the reference setup. Although frame retention was stable across the conditions tested here, opioids alter how the paw is placed independently of load [10], which warrants consideration when comparing the index across compounds. A related constraint is that adapting the bundled pose network to recordings that differ substantially from the reference condition currently requires working with DeepLabCut directly, which reintroduces the computational barrier our platform is intended to lower. Finally, all mice were C57BL/6J. Because the feature set includes pixel brightness and paw contour, it is unclear how well the models will perform in mice with different coat colors.

Taken together, our results show that a standardized, low-cost apparatus combined with bundled models can provide frame-level scoring of nocifensive, pruritic and withdrawal behaviors, together with continuous paw contour metrics and measures of behavioral sequencing, from a single recording. Because the enclosure, acquisition software, pose network and classifiers are released together as one open-source platform, PixelPaws provides a low-cost alternative to commercial systems that returns multiple behavioral and paw-level readouts from every recording rather than a single scored endpoint. Scoring every recording with the same models also makes larger multi-behavior designs practical and removes observer drift from comparisons across cohorts and, in principle, across laboratories.

## Acknowledgments

This work was supported by the National Institutes of Health [grant numbers K99DA056691 and P30DK056341 (Nutrition Obesity Research Center) to R.A.S.; R01DA064355 to R.W.G.; RM1NS135283 and R01DA062509 to M.C.C.]. The authors have no conflicts of interest to declare.

## Data and code availability

PixelPaws, including the analysis interface, the apparatus design files and STLs, the bundled pretrained pose network and the reference behavior classifiers, is released under a CC BY-NC 4.0 license at https://github.com/rslivicki/PixelPaws, and every figure is produced by a script deposited with the code. Raw video and per-frame prediction files are available upon reasonable request to the corresponding author.

## Declaration of generative AI and AI-assisted technologies in the manuscript preparation process

During the preparation of this work the authors used Claude (Anthropic) to assist with manuscript drafting, editing, organization, reference formatting, and figure code. After using this tool, the authors reviewed and edited the content as needed and take full responsibility for the content of the published article.

## Supplementary Material

## Supplementary Methods

Behavioral sequencing: residual construction, display rules and validation

### Residual scale

Pearson residuals were used rather than fold-enrichment because the ratio is unstable wherever the expected count is small, and which cells are small is set by how often each state occurred. On synthetic cohorts with no sequencing structure in either group but differing base rates, fold-enrichment reported a significant difference in 12 of 12 runs. The residuals are not z-scores: the margins were estimated rather than known, the cells are correlated, and the tables were sparse, with a median expected off-diagonal count of 0.23 in the withdrawal cohort. Normality was not required, because the inference is a permutation test.

### Subsampling

Each mouse was subsampled to the cohort’s lowest transition total, drawn 100 times and averaged, so no result depended on a single draw. The 100-event inclusion floor is a pragmatic choice rather than a published standard; it was set before the cohorts were analyzed and applied identically to all of them.

### Permutation

Spaces were enumerated exhaustively and p was the fraction of assignments, the observed one included, with a pseudo-F at least as large as observed: 2⁹ = 512 sign-flip assignments for the nine-animal paired design, and at most 6,435 group assignments for the unpaired cohorts. Because relabeling the two groups is itself an assignment, every assignment in an equal-group design has a mirror with identical pseudo-F, so attainable p values are multiples of 2 divided by the space size. The smallest attainable value is therefore 2/512 = 0.0039 for the paired design and 2/3,432 = 0.0006 for a 7 versus 7 contrast; the naloxone and nerve-injury results sit at these floors.

### Calibration

Animals from a single condition were split into two arbitrary groups, half against the rest, and the full pipeline run, including subsampling, residual construction, distance calculation and permutation testing, so the null was true by construction while sparsity, dispersion and transition totals were unchanged. This was repeated 400 times for each condition arm with a fixed random seed. The false-positive rate at α = 0.05 was 0.045 (18/400; 95% Wilson interval 0.029 to 0.070) in the post-naloxone arm, where the tables are sparsest, and 0.037 (15/400) in the pre-naloxone arm; 0.030 (12/400) and 0.040 (16/400) in the vehicle and oxycodone arms of the nerve-injury cohort; 0.033 (13/400) and 0.028 (11/400) in the saline and chloroquine arms; and 0.000 (0/400; upper bound 0.010) in both arms of the formalin and rimonabant cohorts.

### Network panels

Edge width gives the share of a state’s exits a transition carries; edge color gives its Pearson residual. Pooled tables were rarefied to the smaller group’s transition total, averaged over 100 random draws, because residual magnitude otherwise tracks how much behavior a group produced rather than how it was ordered: in the withdrawal cohort the pooled totals were 2,025 before naloxone against 2,900 during it, which would have inflated every post-naloxone residual by a factor of 1.20. Difference panels draw changes exceeding four Pearson-residual units, a display threshold rather than a significance threshold, applied identically to every cohort; at thresholds of 2, 3, 4, 5, 6 and 8 units the withdrawal cohort showed 15, 12, 8, 7, 6 and 4 of 20 assessable routes. A transition was assessable only where both groups had at least 12 exits from the source state and the route carried at least 6% of them in one group, referred to as the assessability floor; states failing it are drawn dashed and unlabelled because their residuals cannot be interpreted, not because nothing happened there. The panels are descriptive, computed from pooled counts without error bars, and no edge-specific test was performed. PERMANOVA at the animal level is the only inferential claim.

### Sensitivity to event boundaries

Post-processing parameters were specified in frames and therefore corresponded to different minimum bout durations across behaviors. Sensitivity was tested by imposing a minimum dwell time before each sequence was built, dissolving runs shorter than the threshold, shortest first, into the longer of their two neighbors; a run at the start or end of a sequence went to its only neighbor, and ties went to the preceding run. Sensitivity to unscored time was tested by excluding transitions that bridged more than a given gap. Minimum dwell times of 0.10 and 0.25 s left every conclusion unchanged in the withdrawal, nerve-injury and formalin cohorts (withdrawal p = 0.0039 throughout, R² 0.32 to 0.35; nerve injury p = 0.0006 to 0.0013, R² 0.60 to 0.68; formalin p = 0.0087 to 0.0281, R² 0.20 to 0.25), as did gap cutoffs down to 0.5 s (withdrawal p = 0.0039, R² 0.25 to 0.32; nerve injury p = 0.0006 at every cutoff; formalin p = 0.026 to 0.033). At dwell times of 0.5 s and above, and with no bridging of unscored time at all, animals fell below the 100-transition floor and the tests lost power by attrition rather than by loss of effect. These are the same data re-analyzed rather than independent replications; the claim is only that the conclusion does not depend on where the event boundaries were drawn.

### Paw contour filtering and the frame gate

Two filters were applied in sequence. Contours smaller than 4 px² were discarded at extraction. A frame gate then admitted a frame to the index only if the contour area of each hind paw fell between 1,500 and 5,000 px², the mean intensity of each hind-paw contour exceeded zero, and the frame was not scored as licking, since the paw leaves the floor during licking and the contact signal is lost. The area band was set from the empirical distribution: correctly segmented hind-paw contours occupy approximately 2,800 px², with an interquartile range of roughly 2,650 to 3,050 px², so 1,500 to 5,000 px² spans about twofold either side of the typical value within a region of interest of 10,000 px². Sessions lacking a pose file were skipped in their entirety, per-frame pose likelihood was not used as a filter, and this gate was applied to every contour analysis reported here.

Frame retention under the gate was 96.7% in vehicle-paw and 86.0% in formalin-injected mice, 86, 85, 93 and 93% across the oxycodone dose series, and 94.4% and 93.0% at spared nerve injury baseline and post-drug.

### PixelPaws supplemental tables

*Every statistics table is recomputed from the caches the figures are drawn from, so a table cannot disagree with the panel it describes. Post-tests against a common control are Holm corrected within each dose family; raw and adjusted P are both given, and the P-value summary reflects the adjusted value. Group values are mean ± SEM*.

**Table S1.**
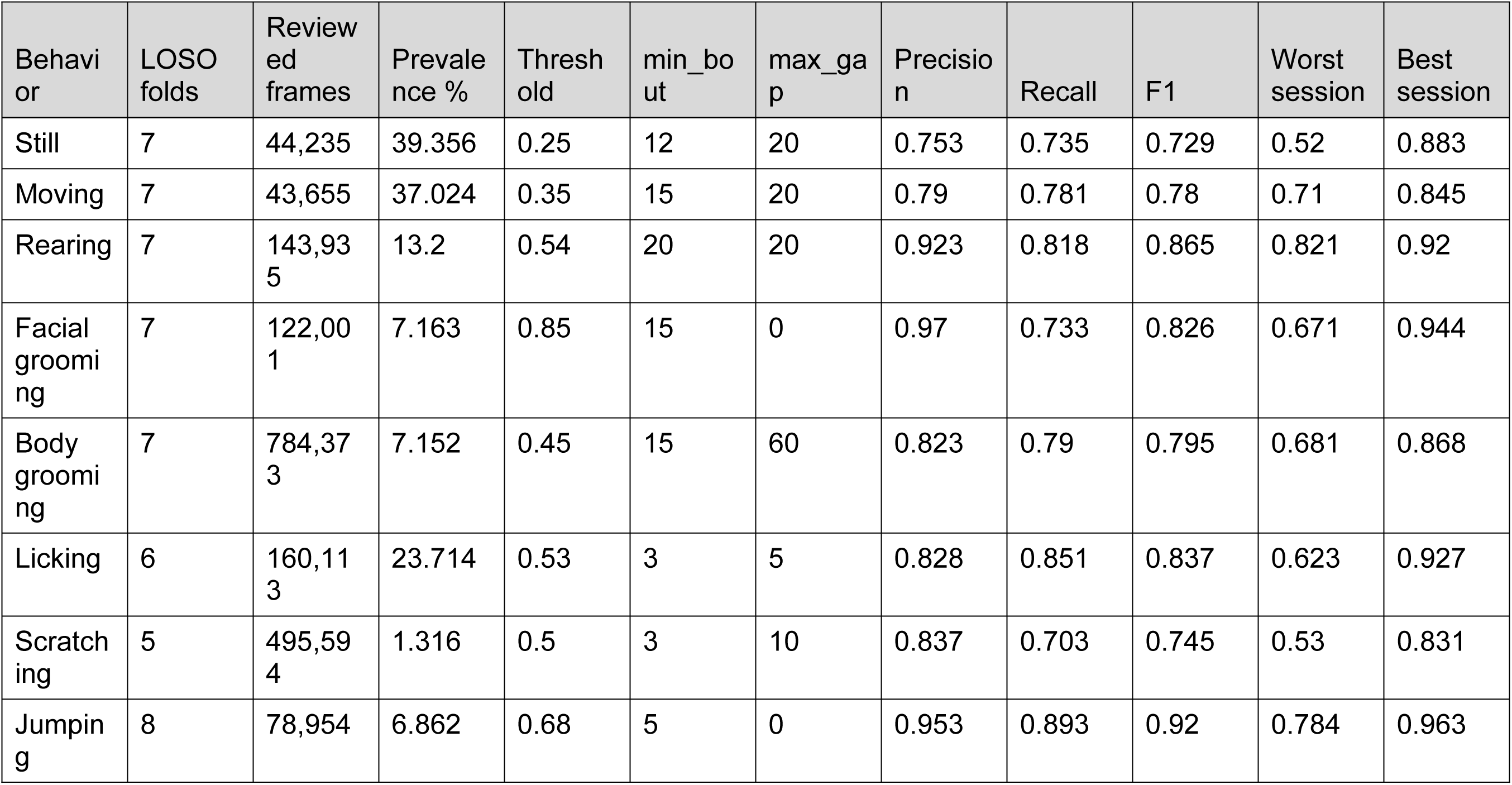
Classifier performance (out-of-fold)

**Table S2.** Recording setup comparison.

| Behavior | Setup 1 (%) | Setup 2 (%) | Difference (pp) | p | n |
| --- | --- | --- | --- | --- | --- |
| Still | 36.08 ± 7.34 | 38.93 ± 6.08 | 2.85 | 0.914 | 6 vs 4 |
| Moving | 24.66 ± 4.94 | 22.81 ± 2.08 | -1.85 | 0.914 | 6 vs 4 |
| Rearing | 9.12 ± 2.57 | 7.93 ± 2.14 | -1.19 | 1 | 6 vs 4 |
| Facial grooming | 10.64 ± 2.10 | 9.05 ± 3.18 | -1.59 | 0.762 | 6 vs 4 |
| Body grooming | 8.94 ± 1.61 | 7.94 ± 2.12 | -1 | 0.61 | 6 vs 4 |
| Licking | 1.47 ± 0.37 | 2.84 ± 0.98 | 1.38 | 0.352 | 6 vs 4 |
| Scratching | 0.78 ± 0.42 | 0.44 ± 0.09 | -0.34 | 0.762 | 6 vs 4 |

**Table S3.** Figure S3, video compression sweep.

| Encoding | n | Duration (s) | File (MB) | Bitrate (Mbit/s) | Smaller than source | State agreement (%) | Pose shift (px) | Paw brightness (%) | Contour ratio (log units) |
| --- | --- | --- | --- | --- | --- | --- | --- | --- | --- |
| source | 4 | 1,847 | 1,869 | 8.1 | - | 100 | 0 | 0 | 0 |
| x264_crf23 | 4 | 180 | 7.78 | 0.35 | 23x | 91.8 | 0.48 | 0.28 | 0.001 |
| x264_crf26 | 4 | 180 | 4.43 | 0.2 | 41x | 89.7 | 0.61 | 0.35 | 0.002 |
| x264_crf28 | 4 | 180 | 3.34 | 0.15 | 55x | 87.8 | 0.7 | 0.43 | 0.002 |
| x264_crf31 | 4 | 180 | 2.36 | 0.1 | 77x | 84.4 | 0.9 | 0.56 | 0.003 |
| x264_crf35 | 4 | 180 | 1.62 | 0.07 | 112x | 80.9 | 1.23 | 0.83 | 0.004 |
| x265_crf28 | 4 | 180 | 3.86 | 0.17 | 47x | 88.9 | 0.59 | 0.33 | 0.002 |

**Table S4.** Oxycodone potency on both readouts.

| Type of test | Comparison | Group 1 vs Group 2 | P-value summary | Statistic, P-value | Effect size |
| --- | --- | --- | --- | --- | --- |
| hyperbolic, top = 100 | licking, ED<sub>50</sub>, phase 1 (0-10 min) | Pantouli 2021: 2.7 (1.4-4.9) mg/kg i.p. |  | ED50 = 0.99 mg/kg | 95% CI 0.56–2.27 |
|  | contour intensity, ED<sub>50</sub>, phase 1 (0-10 min) | fit extrapolates; see reversal below |  | ED50 = 5.39 mg/kg | 95% CI 2.00–27.02 |
| observed | contour intensity, reversal at 1 mg/kg, phase 1 (0-10 min) | 0.908 vs 0.902 (vehicle) HL/HR |  | 6% back to symmetry | - |
|  | contour intensity, reversal at 3 mg/kg, phase 1 (0-10 min) | 0.938 vs 0.902 (vehicle) HL/HR |  | 37% back to symmetry | - |
|  | contour intensity, reversal at 10 mg/kg, phase 1 (0-10 min) | 0.969 vs 0.902 (vehicle) HL/HR |  | 69% back to symmetry | - |
| hyperbolic, top = 100 | licking, ED<sub>50</sub>, phase 2 (10-60 min) | Pantouli 2021: 3.4 (1.9-5.8) mg/kg i.p. |  | ED50 = 3.27 mg/kg | 95% CI 1.56–8.50 |
|  | contour intensity, ED<sub>50</sub>, phase 2 (10-60 min) | fit extrapolates; see reversal below |  | ED50 = 15.28 mg/kg | 95% CI 3.00–500.00 |
| observed | contour intensity, reversal at 1 mg/kg, phase 2 (10-60 min) | 0.903 vs 0.915 (vehicle) HL/HR |  | -14% back to symmetry | - |
|  | contour intensity, reversal at 3 mg/kg, phase 2 (10-60 min) | 0.934 vs 0.915 (vehicle) HL/HR |  | 22% back to symmetry | - |
|  | contour intensity, reversal at 10 mg/kg, phase 2 (10-60 min) | 0.950 vs 0.915 (vehicle) HL/HR |  | 41% back to symmetry | - |

**Table S5.** Statistics from Figure S8, rimonabant dose-response on scratching.

| Type of test | Comparison | Group 1 vs Group 2 | P-value summary | Statistic, P-value | Effect size |
| --- | --- | --- | --- | --- | --- |
| one-way ANOVA | scratching bouts, 60 min, effect of dose | - | ** | F(3, 23) = 5.825, P=0.0041 | $\eta^2 = 0.432$ |
| Welch t | scratching bouts, 60 min, 1 mg/kg vs vehicle | 51.7 ± 9.59 vs 11.9 ± 7.48 bouts | * | t(11.3) = 3.277, P=0.0071 (Holm 0.0162) | Hedges g = 1.75 |
|  | scratching bouts, 60 min, 3 mg/kg vs vehicle | 76.4 ± 15.8 vs 11.9 ± 7.48 bouts | * | t(8.6) = 3.695, P=0.0054 (Holm 0.0162) | Hedges g = 1.98 |
|  | scratching bouts, 60 min, 10 mg/kg vs vehicle | 97 ± 24.8 vs 11.9 ± 7.48 bouts | * | t(5.9) = 3.289, P=0.0170 (Holm 0.0170) | Hedges g = 1.96 |

**Figure S1.**
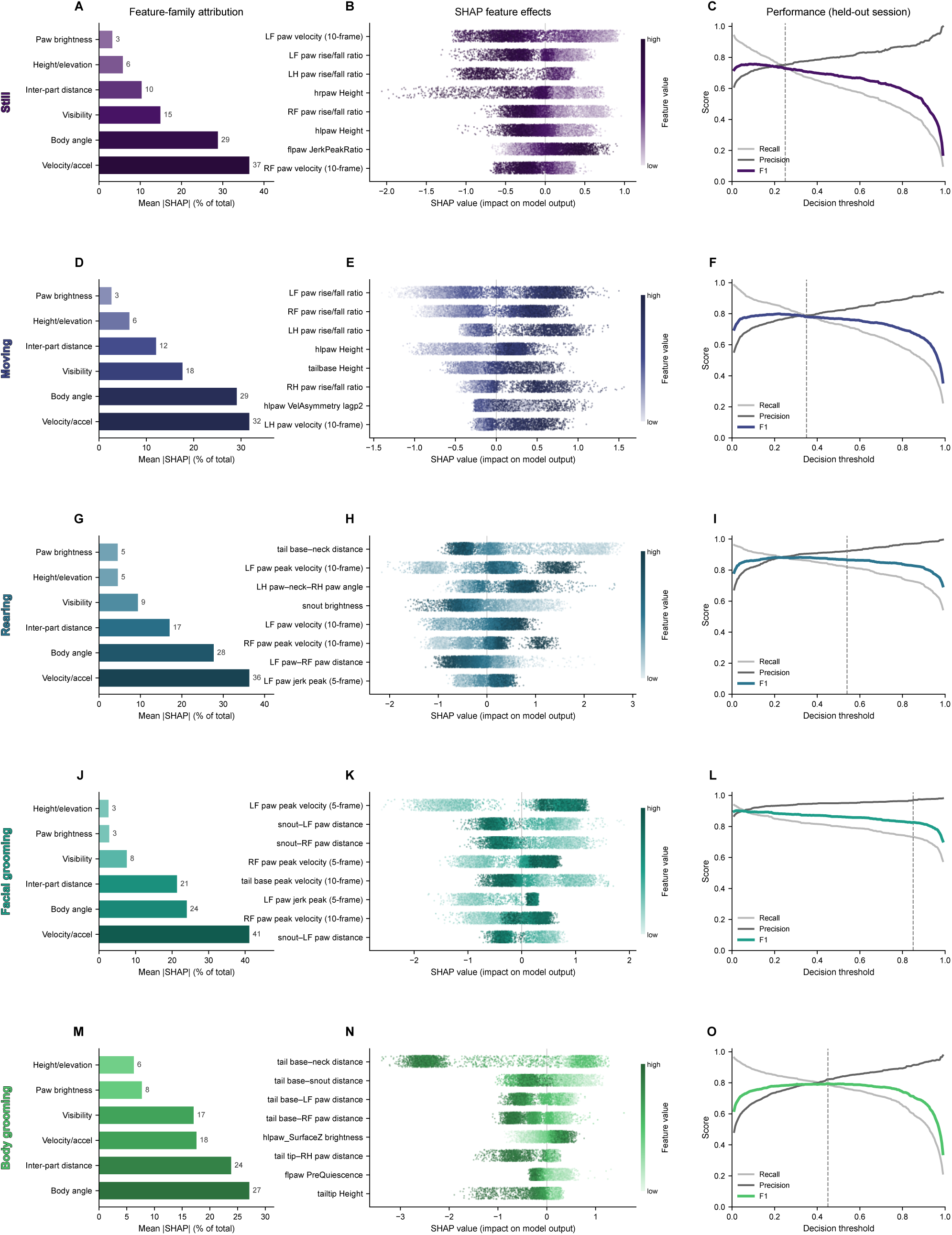
Feature attribution and threshold performance for the five remaining classifiers. Feature-family attribution, expressed as the share of total absolute SHAP value, is shown for stillness (A), moving (D), rearing (G), facial grooming (J) and body grooming (M), with SHAP values for the strongest individual features by beeswarm (B,E,H,K,N) and precision, recall and F1 across the decision threshold (C,F,I,L,O). Every value shown is out of fold, leave-one-session-out across seven annotated sessions, with each classifier’s bout post-processing applied. The dashed line marks the threshold the classifier ships with. Body grooming is shown for a rebuilt classifier that excludes unreviewed frames the shipped one trained on as negatives. Per-behavior F1, between-session spread and frame counts are in Table S1. Data are represented as the mean across held-out sessions. N = 7 sessions.

**Figure S2.**
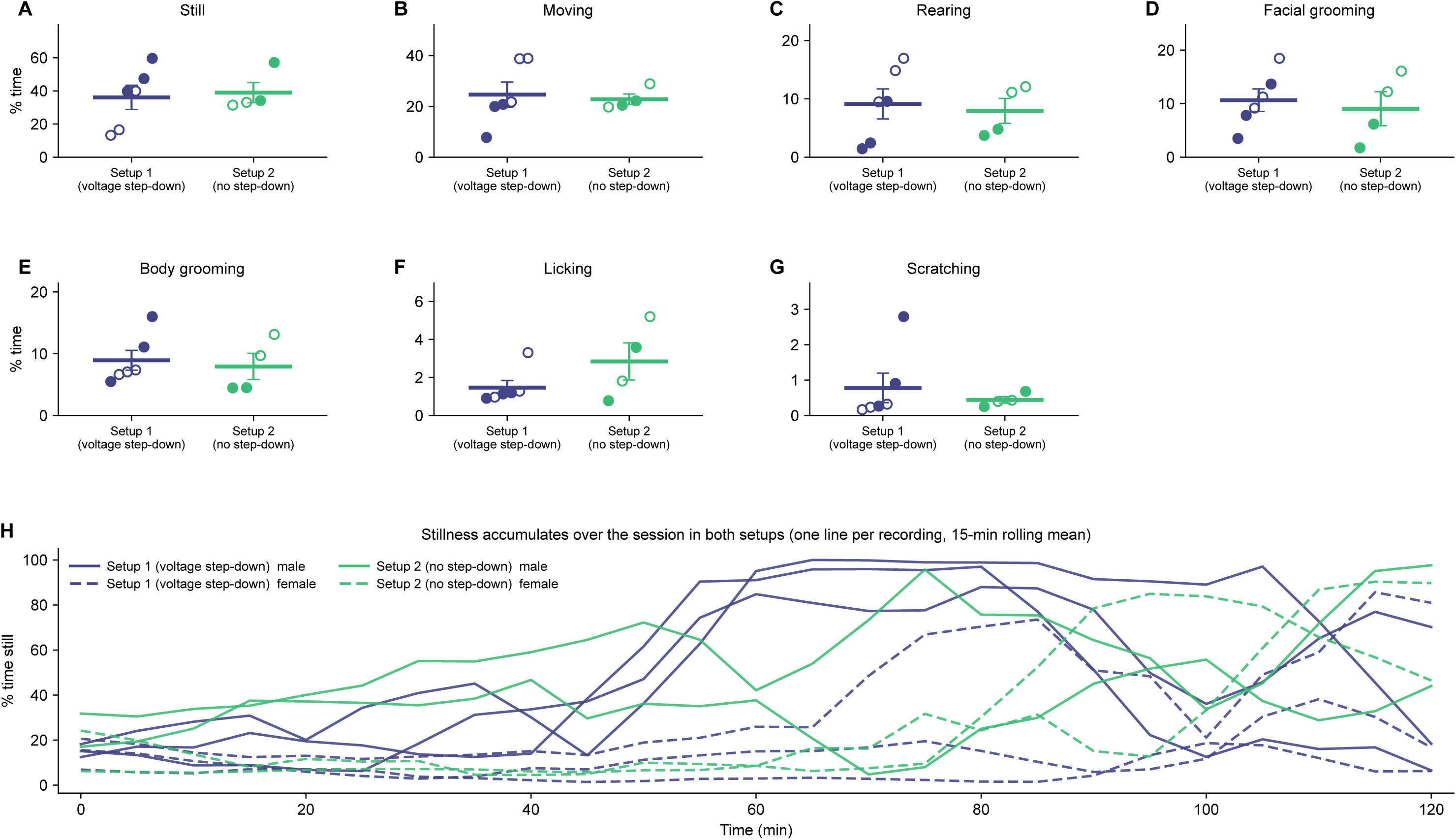
Classifier output does not differ across two recording setups. Ten naive mice were recorded, one session each, each in one of two individual behavioral chambers and scored with the classifiers used throughout this work. Setup 1 used the voltage step-down; Setup 2 did not. Whole-session percent time is shown for stillness, moving, rearing, facial grooming, body grooming, licking and scratching, with the sexes pooled (A-G), and the stillness time course per recording over 120 min (H). No behavior differed between setups. Filled symbols denote male and open symbols female animals. Mann-Whitney test. Data are expressed as group means +/- SEM with individual recordings overlaid. N = 6 recordings for Setup 1 and 4 for Setup 2.

**Figure S3.**
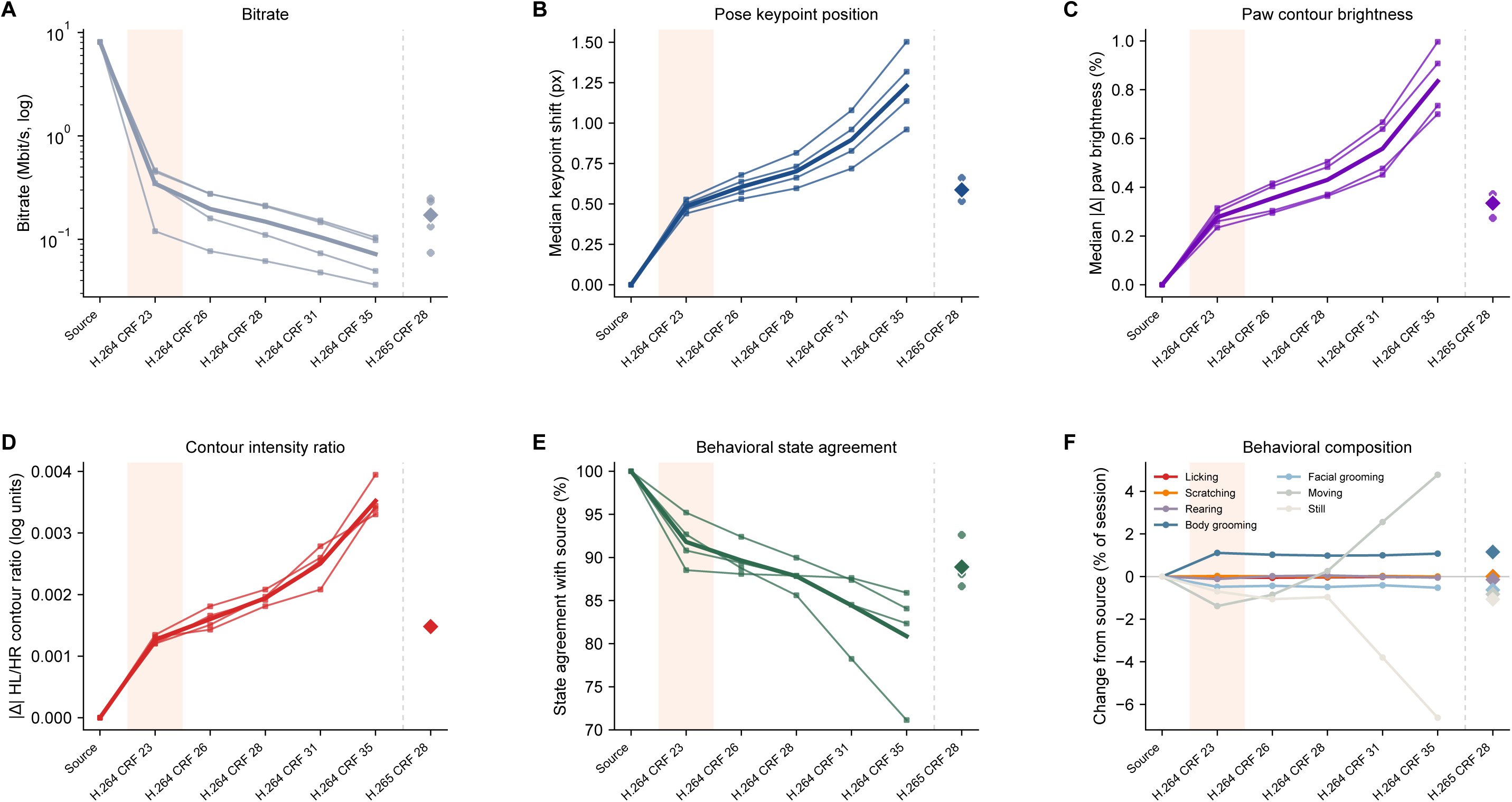
Pose, paw contour, and classifier readouts across video compression rates. A 3-min segment (minutes 5-8) of one session from each of four mice was re-encoded from the source recording, the hardware-encoded H.264 file written at acquisition (about 8 Mbit/s), at five x264 rates and one x265 rate, and each encode was run through the full pipeline (pose estimation, feature extraction and classification) independently of the others. Deviation from the source recording is shown for bitrate (A), pose keypoint position (B), paw contour brightness (C), the hind-paw contour intensity ratio that carries Figures 3-5 (D), frame-by-frame agreement of the behavioral state call (E), and the change in whole-session behavioral composition (F). The x265 encode is a different codec rather than a step on the x264 rate ramp, so it is drawn as a detached diamond beyond the dashed rule. The shaded column marks CRF 23, the rate the preprocessing step applies by default and therefore the rate every analysis in this manuscript was computed on. Data are represented as the mean across animals. N = 4 animals, one segment each.

**Figure S4.**
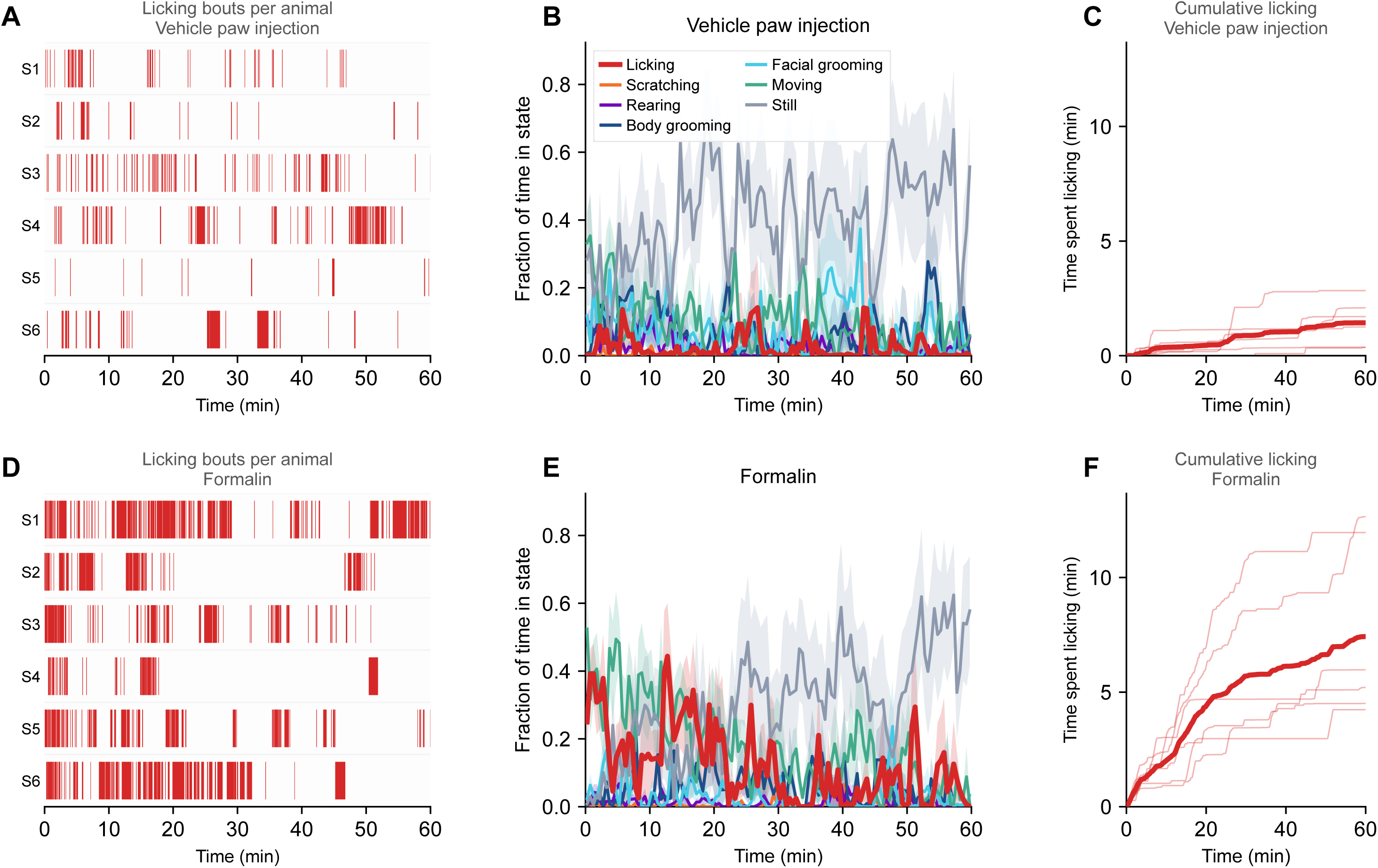
Formalin elevates licking without shifting the rest of the repertoire. Per-animal raster of licking bouts (A,D), fraction of time in each state in non-overlapping 30 s bins (B,E), and cumulative licking time (C,F). Rows: A-C vehicle paw injection; D-F formalin. Animals are numbered S1..Sn; all panels share the same time span. Licking is the readout and is drawn in red. Frames are scored by all seven classifiers and each frame is assigned to a single state by a fixed priority order (Methods 2.7), so cumulative time in the readout can differ from the single-classifier figures, which score each behavior independently. Licking was elevated in formalin-injected animals across the session, while the remaining behaviors were comparable between groups. The state traces (B,E) show mean +/- SEM; rasters and cumulative curves show individual animals, with the group mean overlaid on the cumulative panels. N = 6 per group. Statistics for the licking contrast are in Table 2.

**Figure S5.**
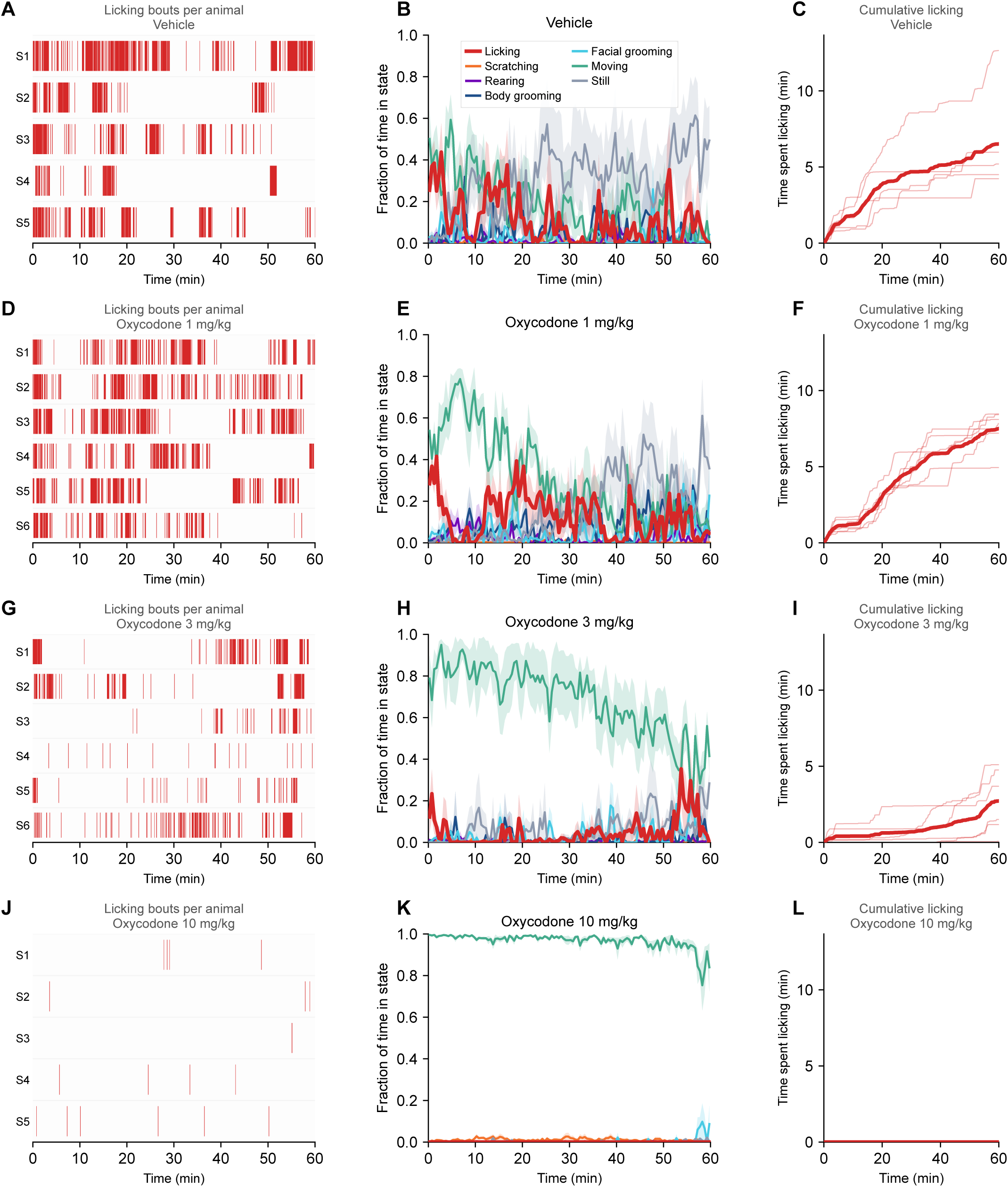
Oxycodone reduces licking dose-dependently and replaces stillness with locomotion. Per-animal raster of licking bouts (A,D,G,J), fraction of time in each state in non-overlapping 30 s bins (B,E,H,K), and cumulative licking time (C,F,I,L). Rows: A-C vehicle; D-F oxycodone 1; G-I oxycodone 3; J-L oxycodone 10 mg/kg. Animals are numbered S1..Sn; all panels share the same time span. Licking is the readout and is drawn in red. Frames are scored by all seven classifiers and each frame is assigned to a single state by a fixed priority order (Methods 2.7), so cumulative time in the readout can differ from the single-classifier figures, which score each behavior independently. Licking fell dose-dependently, and time otherwise spent still was replaced by locomotion. The state traces (B,E,H,K) show mean +/- SEM; rasters and cumulative curves show individual animals, with the group mean overlaid on the cumulative panels. N = 5-6 per group.

**Figure S6.**
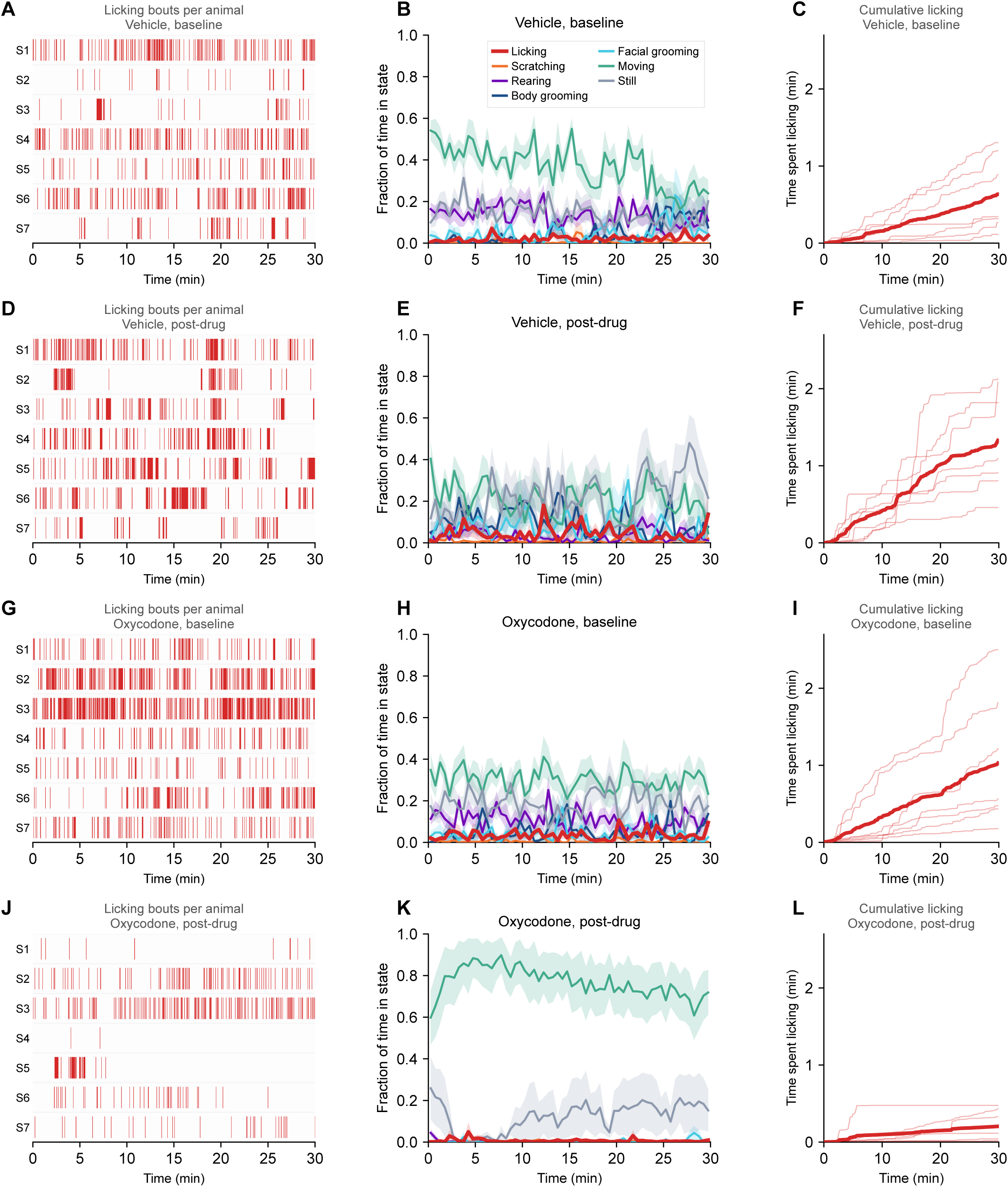
Oxycodone reduces licking and raises locomotion after nerve injury. Per-animal raster of licking bouts (A,D,G,J), fraction of time in each state in non-overlapping 30 s bins (B,E,H,K), and cumulative licking time (C,F,I,L). Rows: A-C vehicle baseline; D-F vehicle post-drug; G-I oxycodone baseline; J-L oxycodone post-drug. Animals are numbered S1..Sn; all panels share the same time span. Licking is the readout and is drawn in red. Frames are scored by all seven classifiers and each frame is assigned to a single state by a fixed priority order (Methods 2.7), so cumulative time in the readout can differ from the single-classifier figures, which score each behavior independently. Oxycodone-treated animals licked less than vehicle-treated animals over the analyzed window and spent more time moving. The state traces (B,E,H,K) show mean +/- SEM; rasters and cumulative curves show individual animals, with the group mean overlaid on the cumulative panels. N = 7 per group, all male. Statistics are in Table 5.

**Figure S7.**
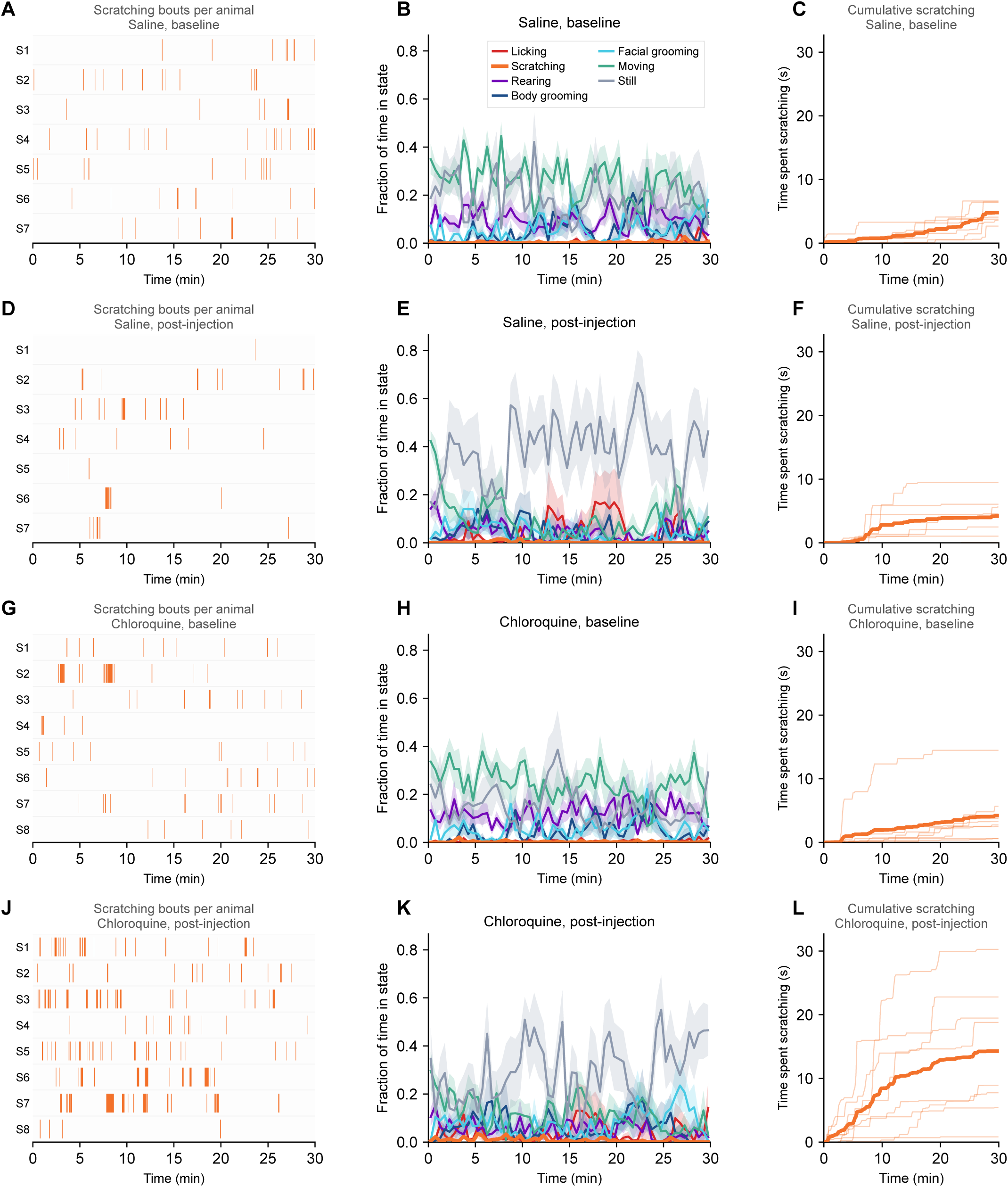
Chloroquine raises scratching without shifting the rest of the repertoire. Per-animal raster of scratching bouts (A,D,G,J), fraction of time in each state in non-overlapping 30 s bins (B,E,H,K), and cumulative scratching time (C,F,I,L). Rows: A-C saline baseline; D-F saline post-injection; G-I chloroquine baseline; J-L chloroquine post-injection. Animals are numbered S1..Sn; all panels share the same time span. Scratching is the readout and is drawn in orange. Frames are scored by all seven classifiers and each frame is assigned to a single state by a fixed priority order (Methods 2.7), so cumulative time in the readout can differ from the single-classifier figures, which score each behavior independently. Scratching rose after chloroquine but not after saline. The state traces (B,E,H,K) show mean +/- SEM; rasters and cumulative curves show individual animals, with the group mean overlaid on the cumulative panels. N = 7-8 per group, all female. Statistics are in Table 6.

**Figure S8.**
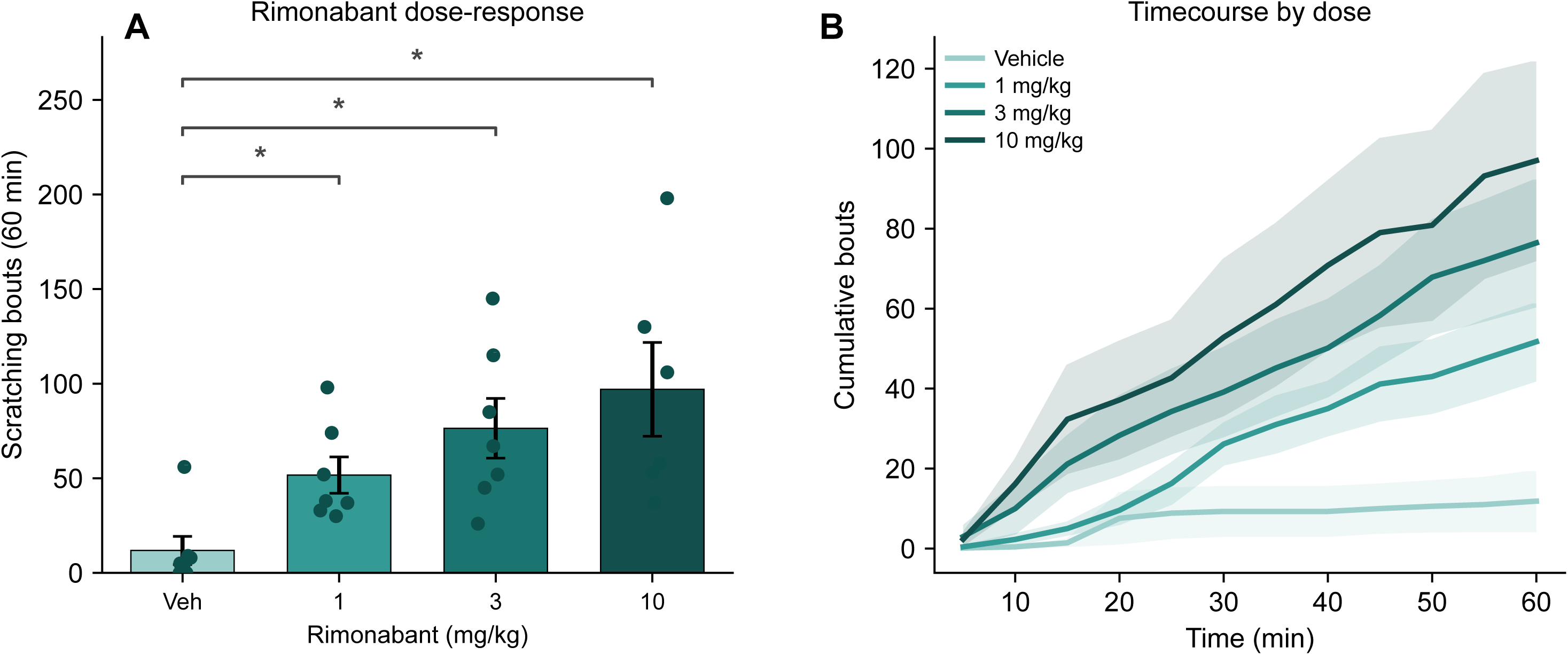
Scratching bouts and cumulative time course across rimonabant doses. The scratching classifier was applied to animals receiving rimonabant (1, 3, 10 mg/kg) or vehicle and scored over 60 min. Total scratching bouts increased dose-dependently (A), with the corresponding cumulative time course (B). ****p<0.0001, ***p<0.001, **p<0.01, *p<0.05 Welch t test versus vehicle (A); one-way ANOVA effect of dose. Post-tests and other statistics are reported in the per-figure statistics table. Data are represented as mean +/- SEM with individual animals overlaid. N = 6-7 per group.

**Figure S9.**
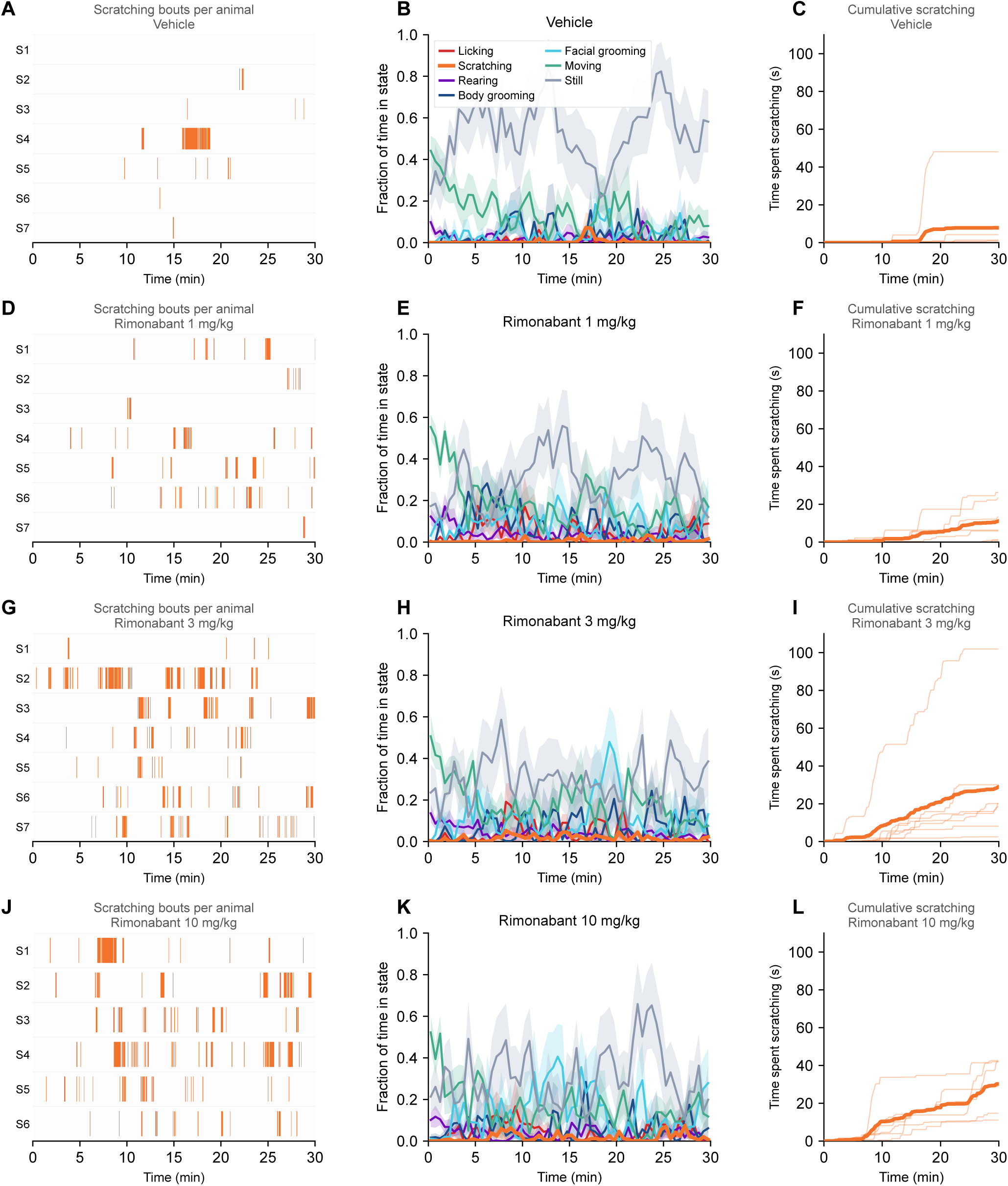
Rimonabant raises scratching dose-dependently without shifting the rest of the repertoire. Per-animal raster of scratching bouts (A,D,G,J), fraction of time in each state in non-overlapping 30 s bins (B,E,H,K), and cumulative scratching time (C,F,I,L). Rows: A-C vehicle; D-F rimonabant 1; G-I rimonabant 3; J-L rimonabant 10 mg/kg. The pooled pre-drug baseline is not drawn as its own row; it appears in the statistics table only. Animals are numbered S1..Sn; all panels share the same time span. Scratching is the readout and is drawn in orange. Frames are scored by all seven classifiers and each frame is assigned to a single state by a fixed priority order (Methods 2.7), so cumulative time in the readout can differ from the single-classifier figures, which score each behavior independently. Panels show the first 30 min of the 60 min assay. Scratching increased with dose, while the remaining behaviors were comparable across dose groups. The state traces (B,E,H,K) show mean +/- SEM; rasters and cumulative curves show individual animals, with the group mean overlaid on the cumulative panels. N = 6-7 per group. This cohort has no matched baseline; see Figure S8.

**Figure S10.**
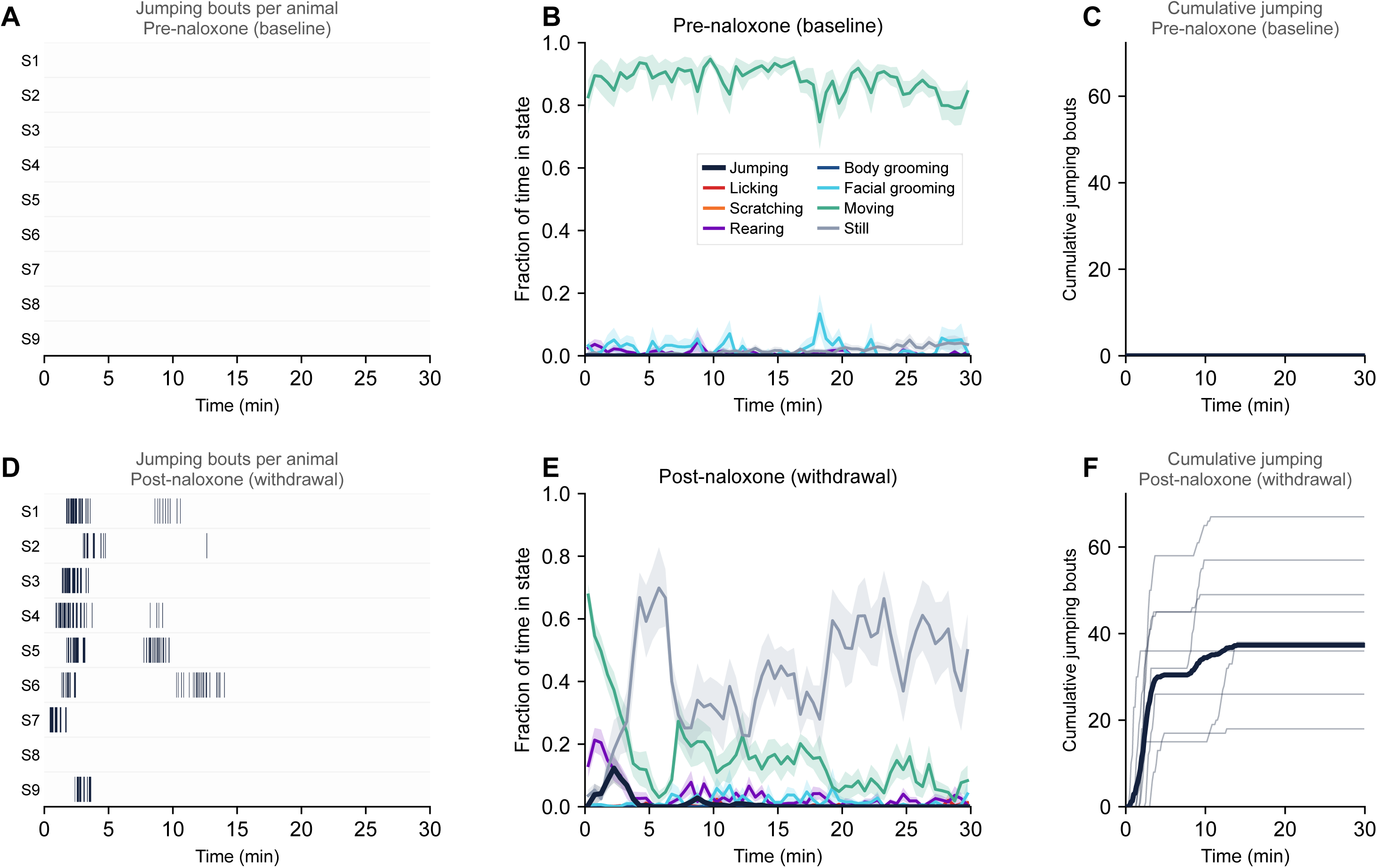
Naloxone reduces locomotion and raises stillness, with jumping appearing against that background. Per-animal raster of jumping bouts (A,D), fraction of time in each state in non-overlapping 30 s bins (B,E), and cumulative jump bouts (C,F). Rows: A-C pre-naloxone; D-F post-naloxone. Animals are numbered S1..Sn; all panels share the same time span. Jumping is the readout and is drawn in purple. Frames are scored by all seven classifiers and each frame is assigned to a single state by a fixed priority order (Methods 2.7), so cumulative time in the readout can differ from the single-classifier figures, which score each behavior independently. Naloxone reduced time spent moving and increased time spent still, with jumping appearing against that background. The state traces (B,E) show mean +/- SEM; rasters and cumulative curves show individual animals, with the group mean overlaid on the cumulative panels. N = 9 female mice per phase, each contributing both. Statistics are in Table 7.

**Figure S11.**
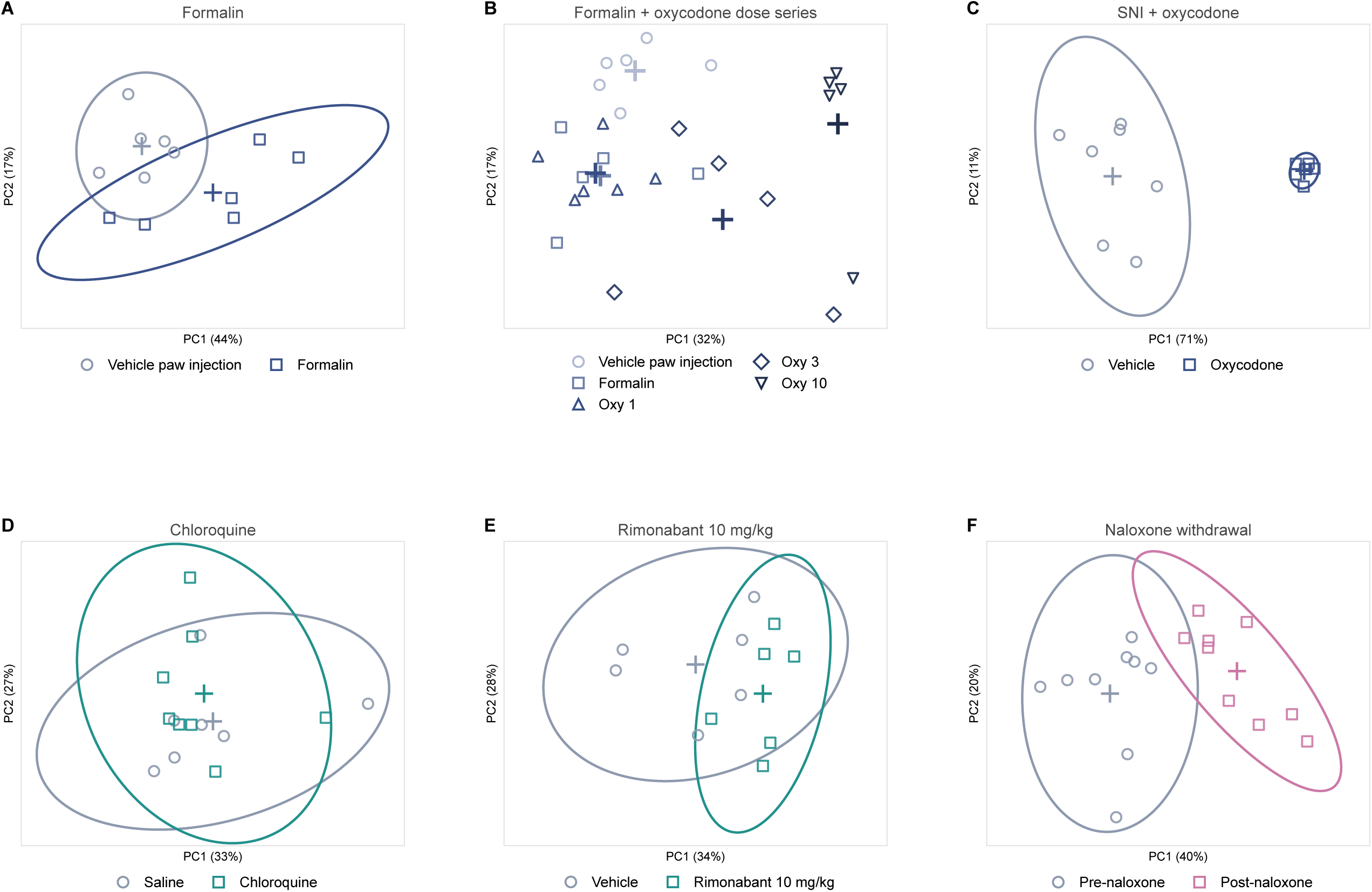
Per-animal behavioral sequencing for every contrast in the battery. Panel D of Figure 8 repeated for each model, in the order the paper introduces them: formalin (A), nerve injury with oxycodone (C), chloroquine (D), rimonabant 10 mg/kg (E) and naloxone-precipitated withdrawal (F). Panel B places the formalin dose series in the same space as a vehicle paw-injection reference. Each point is one animal and the cross is the group mean; control conditions are open circles and treated conditions open squares. In the two-group panels the surrounding line is a 95% normal-theory ellipse describing the spread of that group, not a confidence region for its mean; panel (B) shows animals and group means without ellipses. PERMANOVA; in the naloxone panel (F) the permutation is restricted within animal, and the other contrasts are unpaired. Post-tests and other statistics are reported in the per-figure statistics table. N = 5-9 per group.

## References

[1] Akiba T, Sano S, Yanase T, Ohta T, Koyama M. Optuna: a next-generation hyperparameter optimization framework. In: Proceedings of the 25th ACM SIGKDD International Conference on Knowledge Discovery and Data Mining. Anchorage: Association for Computing Machinery, 2019:2623-2631. doi:10.1145/3292500.3330701.

[2] Anderson MJ. Distance-based tests for homogeneity of multivariate dispersions. Biometrics 2006;62:245–253. doi:10.1111/j.1541-0420.2005.00440.x.

[3] Anderson MJ, Walsh DCI, Clarke KR, Gorley RN, Guerra-Castro E. Some solutions to the multivariate Behrens-Fisher problem for dissimilarity-based analyses. Aust N Z J Stat 2017;59:57–79.

[4] Barkai O, Zhang B, Lenfers Turnes B, Arab M, Yarmolinsky DA, Zhang Z, Barrett LB, Woolf CJ. A machine learning tool with light-based image analysis for automatic classification of 3D pain behaviors. Cell Rep Methods 2025;5:101145. doi:10.1016/j.crmeth.2025.101145.

[5] BioSyft. Automated behavioral analytics for preclinical research [Internet]. Available from: https://biosyft.io/ [accessed 27 August 2026].

[6] BlackBox Bio. Blackbox One, Blackbox Two and palmReader [Internet]. Available from: https://blackboxbio.com/ [accessed 27 August 2026].

[7] Bohnslav JP, Wimalasena NK, Clausing KJ, Dai YY, Yarmolinsky DA, Cruz T, Kashlan AD, Chiappe ME, Orefice LL, Woolf CJ, Harvey CD. DeepEthogram, a machine learning pipeline for supervised behavior classification from raw pixels. eLife 2021;10:e63377.

[8] Bravo IM, Bluitt MN, McElligott ZA. Examining opioid withdrawal scoring and adaptation of global scoring systems to male and female C57BL/6J mice. bioRxiv 2021.10.11.463944 [preprint]. doi:10.1101/2021.10.11.463944.

[9] Bravo IM, Luster BR, Flanigan ME, Perez PJ, Cogan ES, Schmidt KT, McElligott ZA. Divergent behavioral responses in protracted opioid withdrawal in male and female C57BL/6J mice. Eur J Neurosci 2020;51:742–754. doi:10.1111/ejn.14580. PMCID PMC7069788.

[10] Brings VE, Payne MA, Gereau RW 4th. Opioids alter paw placement during walking, confounding assessment of analgesic efficacy in a postsurgical pain model in mice. Pain Rep 2022;7:e1035. doi:10.1097/PR9.0000000000001035. PMCID PMC9416758.

[11] Chen T, Guestrin C. XGBoost: a scalable tree boosting system. In: Proceedings of the 22nd ACM SIGKDD International Conference on Knowledge Discovery and Data Mining. San Francisco: Association for Computing Machinery, 2016:785-794. doi:10.1145/2939672.2939785.

[12] Contreras KM, Buzzi B, Vaughn J, Caillaud M, Altarifi AA, Olszewski E, Walentiny DM, Beardsley PM, Damaj MI. Characterization and validation of a spontaneous acute and protracted oxycodone withdrawal model in male and female mice. Pharmacol Biochem Behav 2024;242:173795. doi:10.1016/j.pbb.2024.173795. PMCID PMC11283946.

[13] Darmani NA, Pandya DK. Involvement of other neurotransmitters in behaviors induced by the cannabinoid CB1 receptor antagonist SR 141716A in naive mice. J Neural Transm 2000;107:931–945.

[14] Decosterd I, Woolf CJ. Spared nerve injury: an animal model of persistent peripheral neuropathic pain. Pain 2000;87:149–158. doi:10.1016/S0304-3959(00)00276-1.

[15] Dubuisson D, Dennis SG. The formalin test: a quantitative study of the analgesic effects of morphine, meperidine, and brain stem stimulation in rats and cats. Pain 1977;4:161–174.

[16] English A, Marcus D, Yadav K, Elkhouly Y, Levy A, Corbit V, Ask M, Scedberg A, Poces-Ball J, Uittenbogaard F, Simons R, Witten I, Zweifel L, Land B, Stella N, Bruchas MR. Behavioral decoding reveals cortical endocannabinoid potentiation during Δ9-THC impairment. bioRxiv 2025.09.26.678874 [preprint]. doi:10.1101/2025.09.26.678874.

[17] Friard O, Gamba M. BORIS: a free, versatile open-source event-logging software for video/audio coding and live observations. Methods Ecol Evol 2016;7:1325–1330. doi:10.1111/2041-210X.12584.

[18] Goodwin NL, Choong JJ, Hwang S, Pitts K, Bloom L, Islam A, Zhang YY, Szelenyi ER, Tong X, Newman EL, Miczek K, Wright HR, McLaughlin RJ, Norville ZC, Eshel N, Heshmati M, Nilsson SRO, Golden SA. Simple Behavioral Analysis (SimBA) as a platform for explainable machine learning in behavioral neuroscience. Nat Neurosci 2024;27:1411–1424. doi:10.1038/s41593-024-01649-9.

[19] Hu Y, Ferrario CR, Maitland AD, Ionides RB, Ghimire A, Watson B, Iwasaki K, White H, Xi Y, Zhou J, Ye B. LabGym: quantification of user-defined animal behaviors using learning-based holistic assessment. Cell Rep Methods 2023;3:100415.

[20] Jacobs GH, Williams GA, Fenwick JA. Influence of cone pigment coexpression on spectral sensitivity and color vision in the mouse. Vision Res 2004;44:1615–1622. doi:10.1016/j.visres.2004.01.016.

[21] Kabra M, Robie AA, Rivera-Alba M, Branson S, Branson K. JAABA: interactive machine learning for automatic annotation of animal behavior. Nat Methods 2013;10:64–67. doi:10.1038/nmeth.2281.

[22] Kapoor S, Narayanan A. Leakage and the reproducibility crisis in machine-learning-based science. Patterns 2023;4:100804. doi:10.1016/j.patter.2023.100804.

[23] Kobayashi K, Matsushita S, Shimizu N, Masuko S, Yamamoto M, Murata T. Automated detection of mouse scratching behaviour using convolutional recurrent neural network. Sci Rep 2021.

[24] Li M, Bao Y, Chen M. Behavioral assessment of pain in rodents: advances from evoked responses to spontaneous states and multimodal approaches. Front Pain Res 2026;7:1739384. doi:10.3389/fpain.2026.1739384.

[25] Lundberg SM, Lee SI. A unified approach to interpreting model predictions. In: Advances in Neural Information Processing Systems 30. 2017. p. 4765-4774.

[26] Luster BR, Cogan ES, Schmidt KT, Pati D, Pina MM, Dange K, McElligott ZA. Inhibitory transmission in the bed nucleus of the stria terminalis in male and female mice following morphine withdrawal. Addict Biol 2020;25:e12748. doi:10.1111/adb.12748.

[27] Mathis A, Mamidanna P, Cury KM, Abe T, Murthy VN, Mathis MW, Bethge M. DeepLabCut: markerless pose estimation of user-defined body parts with deep learning. Nat Neurosci 2018;21:1281–1289. doi:10.1038/s41593-018-0209-y.

[28] O’Brien DE, Brenner DS, Gutmann DH, Gereau RW. Assessment of pain and itch behavior in a mouse model of neurofibromatosis type 1. J Pain 2013;14:628–637.

[29] Oswell CS, Rogers SA, James JG, McCall NM, Hsu AI, Salimando GJ, Mahmood M, Wooldridge LM, Wachira M, Jo AY, Sandoval Ortega RA, Wojick JA, Beattie K, Farinas SA, Chehimi SN, Rodrigues A, Wu JWK, Ejoh LL, Kimmey BA, Lo E, Azouz G, Vasquez JJ, Banghart MR, Beier KT, Creasy KT, Crist RC, Ramakrishnan C, Reiner BC, Deisseroth K, Yttri EA, Corder G. Mimicking opioid analgesia in cortical pain circuits. Nature 2026;649:938–947. doi:10.1038/s41586-025-09908-w.

[30] Otsu N. A threshold selection method from gray-level histograms. IEEE Trans Syst Man Cybern 1979;9:62–66. doi:10.1109/TSMC.1979.4310076.

[31] Pantouli F, Grim TW, Schmid CL, Acevedo-Canabal A, Kennedy NM, Cameron MD, Bannister TD, Bohn LM. Comparison of morphine, oxycodone and the biased MOR agonist SR-17018 for tolerance and efficacy in mouse models of pain. Neuropharmacology 2021;185:108439.

[32] Pereira TD, Tabris N, Matsliah A, Turner DM, Li J, Ravindranath S, Papadoyannis ES, Normand E, Deutsch DS, Wang ZY, McKenzie-Smith GC, Mitelut CC, Castro MD, D’Uva J, Kislin M, Sanes DH, Kocher SD, Wang SSH, Falkner AL, Shaevitz JW, Murthy M. SLEAP: a deep learning system for multi-animal pose tracking. Nat Methods 2022;19:486–495. doi:10.1038/s41592-022-01426-1.

[33] Samineni VK, Grajales-Reyes JG, Grajales-Reyes GE, Tycksen E, Copits BA, Pedersen C, Ankudey ES, Sackey JN, Sewell SB, Bruchas MR, Gereau RW. Cellular, circuit and transcriptional framework for modulation of itch in the central amygdala. eLife 2021;10:e68130. doi:10.7554/eLife.68130.

[34] Schlosburg JE, O’Neal ST, Conrad DH, Lichtman AH. CB1 receptors mediate rimonabant-induced pruritic responses in mice: investigation of locus of action. Psychopharmacology (Berl) 2011;216:323–331. doi:10.1007/s00213-011-2224-5.

[35] Severino AL, Mittal N, Hakimian JK, Velarde N, Minasyan A, Albert R, Torres C, Romaneschi N, Johnston C, Tiwari S, Lee AS, Taylor AM, Gaveriaux-Ruff C, Kieffer BL, Evans CJ, Cahill CM, Walwyn WM. Mu-opioid receptors on distinct neuronal populations mediate different aspects of opioid reward-related behaviors. eNeuro 2020;7(4).

[36] Shepherd AJ, Mohapatra DP. Pharmacological validation of voluntary gait and mechanical sensitivity assays associated with inflammatory and neuropathic pain in mice. Neuropharmacology 2017;130:18–29. doi:10.1016/j.neuropharm.2017.11.036.

[37] Slivicki RA, Yi J, Brings VE, Huynh PN, Gereau RW 4th. The cannabinoid agonist CB-13 produces peripherally mediated analgesia in mice but elicits tolerance and signs of central nervous system activity with repeated dosing. Pain 2022;163:1603-1621. doi:10.1097/j.pain.0000000000002550.

[38] Tjolsen A, Berge OG, Hunskaar S, Rosland JH, Hole K. The formalin test: an evaluation of the method. Pain 1992;51:5–17. doi:10.1016/0304-3959(92)90003-T.

[39] Valtcheva MV, Davidson S, Zhao C, Leitges M, Gereau RW. Protein kinase Cδ mediates histamine-evoked itch and responses in pruriceptors. Mol Pain 2015;11:1.

[40] von Ziegler L, Sturman O, Bohacek J. Big behavior: challenges and opportunities in a new era of deep behavior profiling. Neuropsychopharmacology 2021;46:33–44. doi:10.1038/s41386-020-0751-7.

[41] Wang J, Karbasi P, Wang L, Meeks JP. A layered, hybrid machine learning analytic workflow for mouse risk assessment behavior. eNeuro 2023;10:ENEURO.0335-22.2022. doi:10.1523/ENEURO.0335-22.2022.

[42] Ye S, Filippova A, Lauer J, Schneider S, Vidal M, Qiu T, Mathis A, Mathis MW. SuperAnimal pretrained pose estimation models for behavioural analysis. Nat Commun 2024;15:5165. doi:10.1038/s41467-024-48792-2.

[43] Zhang Z, Roberson DP, Kotoda M, Boivin B, Bohnslav JP, González-Cano R, Yarmolinsky DA, Turnes BL, Wimalasena NK, Neufeld SQ, Barrett LB, Quintão NLM, Fattori V, Taub DG, Wiltschko AB, Andrews NA, Harvey CD, Datta SR, Woolf CJ. Automated preclinical detection of mechanical pain hypersensitivity and analgesia. Pain 2022;163:2326–2336. doi:10.1097/j.pain.0000000000002680.

